# Engineered Enzymes that Retain and Regenerate their Cofactors Enable Continuous-Flow Biocatalysis

**DOI:** 10.1101/568972

**Authors:** Carol J. Hartley, Charlotte C. Williams, Judith A. Scoble, Quentin I. Churches, Andrea North, Nigel G. French, Tom Nebl, Greg Coia, Andrew C. Warden, Greg Simpson, Andrew R. Frazer, Chantel Nixon Jensen, Nicholas J. Turner, Colin Scott

## Introduction

**Biocatalysis is used for many chemical syntheses due to its high catalytic rates, specificities and operation under ambient conditions ^1, 2^. Continuous-flow chemistry offers advantages to biocatalysis, avoiding process issues caused by substrate/product inhibition, equilibrium controlled limitations on yield and allosteric control ^3^. Modular continuous-flow biochemistry would also allow the flexible assembly of different complex multistep reactions ^3–6^. Here we tackle some technical challenges that currently prohibit the wide-spread use of continuous flow biocatalysis; cofactor immobilization and site-specific immobilization. We provide the first example of enzymes engineered to retain and recycle their cofactors, and the use of these enzymes in continuous production of chiral pharmaceutical intermediates.**

Enzyme immobilization for continuous-flow applications has been studied for some time; indeed, a number of industrial processes are currently based on such technologies ^4, 6–10^. However, such processes have generally been limited to cofactor-independent enzymes, such as esterases. Cofactors, such as nicotinamide adenine dinucleotide (NAD^+^) and adenosine triphosphate (ATP), are used stoichiometrically unless recycled, typically by a second enzyme: without recycling, cofactors become prohibitively expensive for most industrial syntheses. As cofactors require diffusion for recycling, they are ill-suited for use in continuous-flow reactors and the lack of a practical engineering solution for the issue has stymied the use of cofactor-dependent enzymes in continuous-flow applications ^4^; although, growing interest in immobilized biocatalysts for cell-free metabolic engineering has led to the development of a variety of enzyme-cofactor-carrier combinations ^5^.

Herein, we propose a novel and generalizable chemo-genetic enzyme engineering approach that enables the fabrication of modular, multistep, biocatalytic, continuous-flow reactors using cofactor-dependent enzymes, thereby extending the utility of biocatalysis for continuous-flow production systems and cell-free metabolic engineering ^11^.

## Results

### Nanomachine design

Our design for a biocatalyst that can retain and recycle its cofactor (a ‘nanomachine’) was inspired by enzymes that retain their substrates *via* covalent attachment during a reaction cascade involving multiple active sites, e.g. phosphopantetheine-dependent synthases ^12–14^ and lipoic acid-dependent dehydrogenases ^15^. In such enzymes, the substrate is delivered from one active site to the next by a flexible ‘swinging arm’ that is covalently attached to the protein. We have adopted a similar strategy, whereby a flexible swinging arm covalently attaches a cofactor to a synthetic, multidomain protein and delivers that cofactor to the different active sites of the fusion protein, allowing its simultaneous use and recycling, while preventing its diffusion (Figure 1).

**Fig. 1.**
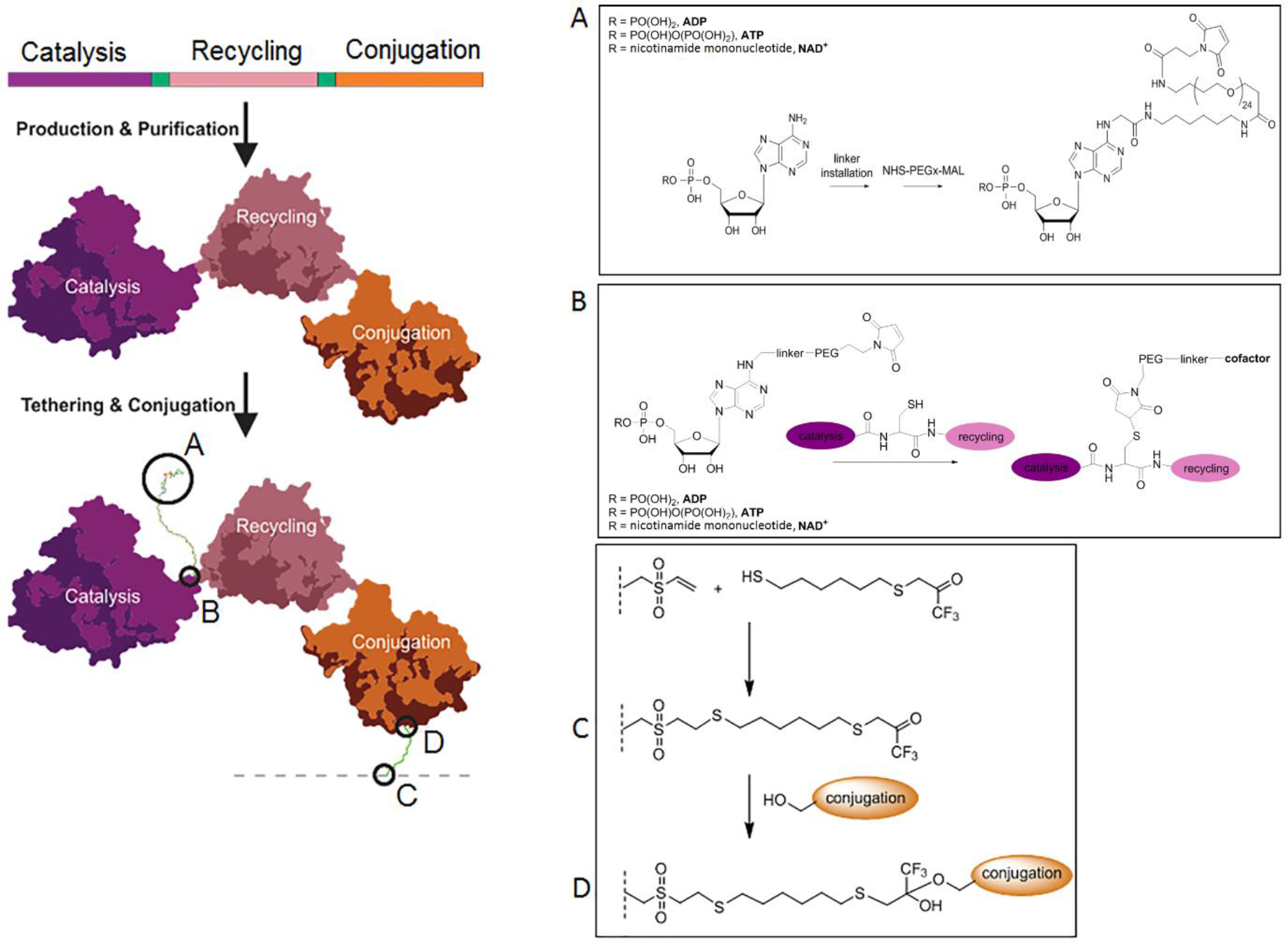
Nanomachine design. Each nanomachine comprises a genetically encoded multi-enzyme fusion protein capable of retaining and recycling a tethered cofactor. The nanomachine contains three protein domains: a cofactor-dependant catalytic enzyme domain (purple), a cofactor-recycling domain (pink) with short amino acid spacer regions between these domains (see Online Methods for details) A cofactor that has been modified by amine activation to allow for linker installation (**A**) is tethered to the protein through maleimide: thiol conjugation *via* a solvent exposed cysteine located in the spacer region between the catalysis and recycling domains (**B**). The esterase conjugation domain (orange) allows immobilization of the nanomachine to a surface by the formation of a covalent bond between a surface attached trifluoroketone (**C**) and the active site serine of the esterase (**D**).

The general design of our nanomachines is shown in Figure 1. Each nanomachine is comprised of three modules: a catalytic module that drives the desired synthesis reaction, a cofactor recycling module that regenerates the cofactor after use, and an immobilization module that allows site-specific, covalent conjugation to an activated surface. The modular design is intended to allow a small number of immobilization and cofactor recycling modules to be used with a wide variety of synthesis modules, thereby enabling diverse synthetic reactions using a relatively small library of core nanomachine components. It is envisaged that multiple nanomachines could be combined in series or in networks to produce a ‘nanofactory’ for the synthesis of complex chiral molecules using multi-enzyme cascade reactions.

In our design, the three modules of each nanomachine were encoded by a single gene for production in *E. coli* as a single protein with a short spacer (2-20 amino acids, Online methods) separating each module. A modified cofactor was designed that could be conjugated to the spacer between the catalytic and cofactor recycling modules. We used a maleimide-functionalized polyethylene glycol (PEG) for the flexible linker to allow movement of the modified cofactor between active sites. *In silico* modelling suggested that a chain length of twenty-four ethylene glycol units was long enough to allow ingress into both active sites. The amino acid spacer that separated the catalytic and cofactor recycling modules contained a single solvent exposed cysteine residue, which provided an accessible thiol group with which to tether the maleimide-functionalized PEGylated cofactor (Figure 1).

### Assembling the nanomachines

For the prototype nanofactory we selected D-fagomine synthesis – coupling three nanomachines in series to convert glycerol (**1**) and 3-aminopropanal (**3**) into a chiral drug precursor (Figure 2). Total synthesis of this anti-diabetic piperidine iminosugar using non-biological catalysts is challenging, due to the complexity conferred by its two stereocenters, and eight possible diastereomers, with even recent advances only resulting in yields up to 65% of each diastereomer ^16^. In our nanofactory (a cascade of nanomachines), glycerol is converted to dihydroxyacetone phosphate (DHAP) (**4**) by regiospecific phosphorylation and oxidation, *via* ATP and NAD^+^-dependent steps, respectively. A subsequent aldolase-catalyzed stereoselective aldol addition with 3-aminopropanal yields (3*S*,4*R*)-amino-3,4-dihydroxy-2-oxyhexyl phosphate (3*S*,4*R*-ADHOP), which can be dephosphorylated with phosphorylase and cyclized to form D-fagomine ^17^ (**6**). For the purposes of purification, we elected to use the carboxybenzyl (Cbz)-protected derivative of 3-aminopropanal, yielding *N-*Cbz-3*S*,4*R*-ADHOP (**5**).

**Figure 2.**
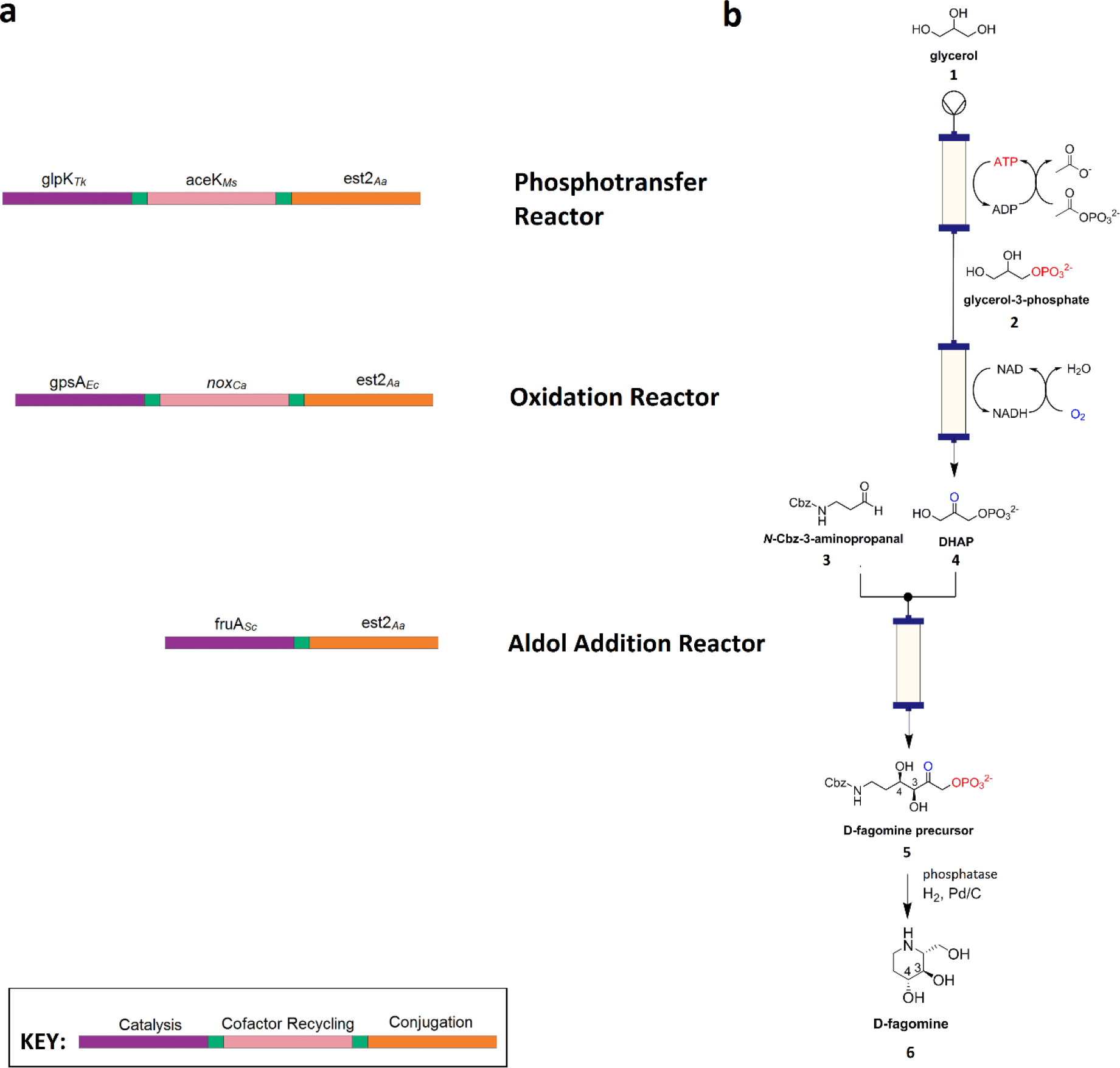
Nanofactory design for the conversion of glycerol to a chiral D-fagomine precursor. **a**, Composition of the three nanomachines each comprising a cofactor-dependant catalytic enzyme domain (purple), a cofactor-recycling domain (pink) and the conjugation domain (orange), with amino acid spacer regions between these domains (green). **b**, Corresponding three part nanofactory and associated biotransformations (phosphotransfer, oxidation and aldol addition) for D-fagomine synthesis. Enzyme name abbreviations are as defined in the text.

Prior comparison of enzymes that could be used for the first two steps of the model synthesis ^18^ had suggested that the most suitable enzymes for glycerol phosphorylation were a *Thermococcus kodakarensis* glycerol kinase (GlpK_Tk_) and a *Mycobacterium smegmatis* acetate kinase (AceK_Ms_). For the NAD^+^-dependent production of DHAP from glycerol-3-phosphate (G3P), *E. coli* glycerol-3-phosphate dehydrogenase (G3PD_Ec_) and the water-forming NADH oxidase from *Clostridium aminovalericum* (NOX_Ca_) were selected. The third reaction step is a cofactor-independent aldolase-catalyzed aldol addition. A monomeric fructose aldolase (FruA) homolog from *Staphylococcus carnosus* was selected from a panel of five potential aldolases. Each of the enzymes incorporated into the nanomachine fusion proteins were selected for their compatibility in batch reactions ^18^, high catalytic rates, relatively simple quaternary structures and thermostability (Supplementary Table 2).

For the conjugation module, we used a serine hydrolase enzyme coupled with a suicide inhibitor (trifluoroketone, TFK) that forms a site-specific and stable covalent bond between the inhibitor and the catalytic serine residue ^19^. Esterase E2 from *Alicyclobacillus acidocaldarius* (E2_Aa_) ^20^ was selected for the serine hydrolase component as a highly stable, soluble, monomeric protein (Supplementary Table 2).

The genes encoding GlpK_Tk_ and AceK_Ms_ were fused, such that GlpK_Tk_ formed the N-terminus of the resultant protein and AceK_Ms_ formed the C-terminus (Fig. 2). The two modules were separated by a nineteen amino acid unstructured amino acid linker, containing a single, solvent-accessible cysteine residue that was used subsequently as the attachment point for the maleimide functionalized PEG_24_-ATP ([GSS]_3_C[GSS]_3_). A similar gene fusion was constructed used G3PD_Ec_ and NOX_Ca_, producing a protein in which G3PD_Ec_ formed the N-terminus and NOX_Ca_ formed the C-terminus. The fused proteins retained or in some cases improved their original catalytic functions (Table 1), albeit some loss of activity (both *K*_M_ and *k*_cat_) was incurred for AceK_Ms_ and G3PD_Ec_. The thermal stability of each of the fused enzymes appeared to be independent of one another i.e. protein unfolding of the component modules of each fusion are independent events (Supplementary Table 2).

**Table 1.** Steady-state kinetic data for the enzymes comprising each nanomachine.

| | $K_M$<br>($\mu\text{M}$) | $K_M$<br>($\mu\text{M}$) | $k_{\text{cat}}$<br>($\text{s}^{-1}$) | $k_{\text{cat}}/K_M$<br>($\text{s}^{-1}\cdot\text{M}^{-1}$) |
| --- | --- | --- | --- | --- |
|  | Cofactor | Substrate |  |  |
| **Glycerol phosphorylation** |  |  |  |  |
| GlpK<sub>TK</sub> | 111 ± 12 | 15 ± 2 | 940 ± 8 | 6.1 × 10<sup>7</sup> |
| GlpK<sub>TK</sub>-AceK<sub>MS</sub> | 123 ± 21 | 15 ± 4 | 1,125 ± 115 | 7.7 × 10<sup>7</sup> |
| GlpK<sub>TK</sub>-AceK<sub>MS</sub>-Est2<sub>Aa</sub> | 115 ± 19 | 16 ± 4 | 1,399 ± 54 | 8.6 × 10<sup>7</sup> |
| GlpK<sub>TK</sub>-ATP<sub>teth</sub>-AceK<sub>MS</sub>-Est2<sub>Aa</sub> | ND | 16 ± 2 | 1,408 ± 95 | 8.8 × 10<sup>7</sup> |
| **Acetate dephosphorylation (ADP phosphorylation)** |  |  |  |  |
| AceK<sub>MS</sub> | 113 ± 9 | 390 ± 8 | 1,103 ± 126 | 2.8 × 10<sup>6</sup> |
| GlpK<sub>TK</sub>-AceK<sub>MS</sub> | 424 ± 35 | 1,400 ± 126 | 759 ± 53 | 5.4 × 10<sup>5</sup> |
| GlpK<sub>TK</sub>-AceK<sub>MS</sub>-Est2<sub>Aa</sub> | 398 ± 29 | 1,197 ± 114 | 1,084 ± 27 | 9.1 × 10<sup>5</sup> |
| GlpK<sub>TK</sub>-ATP<sub>teth</sub>-AceK<sub>MS</sub>-Est2<sub>Aa</sub> | ND | ND | ND | ND |
| **Glycerol-3-phosphate oxidation** |  |  |  |  |
| G3PD<sub>Ec</sub> | 158 ± 24 | 59 ± 4 | 85 ± 11 | 1.4 × 10<sup>6</sup> |
| G3PD<sub>Ec</sub>-NOX<sub>Ca</sub> | 176 ± 12 | 369 ± 17 | 7 ± 0.7 | 1.8 × 10<sup>4</sup> |
| G3PD<sub>Ec</sub>-NOX<sub>Ca</sub>-Est2<sub>Aa</sub> | 164 ± 10 | 659 ± 47 | 7 ± 0.6 | 1.1 × 10<sup>4</sup> |
| G3PD<sub>Ec</sub>-NAD<sub>teth</sub>-NOX<sub>Ca</sub>-Est2<sub>Aa</sub> | ND | 659 ± 47 | 9 ± 0.6 | 1.4 × 10<sup>4</sup> |
| **Oxygen reduction (NADH oxidation)** |  |  |  |  |
| Nox<sub>Ca</sub> | 258 ± 21 | ND | 1,252 ± 182 | 4.9 × 10<sup>6</sup> |
| G3PD<sub>Ec</sub>-NOX<sub>Ca</sub> | 276 ± 9 | ND | 1,714 ± 252 | 6.2 × 10<sup>6</sup> |
| G3PD<sub>Ec</sub>-NOX<sub>Ca</sub>-Est2<sub>Aa</sub> | 266 ± 15 | ND | 1,224 ± 114 | 4.6 × 10<sup>6</sup> |
| G3PD<sub>Ec</sub>-NAD<sub>teth</sub>-NOX<sub>Ca</sub>-Est2<sub>Aa</sub> | ND | ND | ND | ND |
| **Aldol addition** |  |  | **DHAP** |  |
| FruA<sub>Sc</sub> | NA | 500 ± 80 | 16 ± 2 | 3.2 × 10<sup>4</sup> |
| FruA<sub>Sc</sub>-Est2<sub>Aa</sub> | NA | 70 ± 11 | 9 ± 1 | 1.3 × 10<sup>5</sup> |
| **Esterase<sup>1</sup>** |  |  |  |  |
| Est2<sub>Aa</sub> | NA | 180 ± 40 | 153 ± 12 | 8.3 × 10<sup>5</sup> |
| GlpK<sub>TK</sub>-AceK<sub>MS</sub>-Est2<sub>Aa</sub> | NA | 165 ± 18 | 149 ± 16 | 8.8 × 10<sup>5</sup> |
| G3PD<sub>Ec</sub>-NOX<sub>Ca</sub>-Est2<sub>Aa</sub> | NA | 152 ± 28 | 159 ± 11 | 9.0 × 10<sup>5</sup> |
| FruA<sub>Sc</sub>-Est2<sub>Aa</sub> | NA | 100 ± 10 | 165 ± 18 | 1.7 × 10<sup>6</sup> |

<sup>1</sup> (*p*-nitrophenyl acetate as substrate). ND- not determined. NA- not applicable.

The gene encoding the conjugation module (E2_Aa_) was fused with both the *glpK*_*Tk*_-*aceK*_*M*_s and *g3pD*_*Ec*_*-nox*_*Ca*_ such that it formed the C-terminus of the encoded proteins (i.e., GlpK_Tk_-AceK_Ms_-E2_Aa_ and G3PD_Ec_-NOX_Ca_-E2_Aa_). Addition of the conjugation module had little effect on the kinetic performance or thermal stabilities of the other modules of each nanomachine (Table 1 & Supplementary Table 1). The conjugation module was highly efficient (86-98% immobilization efficiency), and the immobilized enzymes retained their activity. This immobilization technique has the potential for broad applicability to other biocatalytic systems.

The cofactors were modified (Online methods) by functionalization of the C6-adenine-amine of ADP (**7**) or NAD^+^ (**12**) to which a modified PEG_24_ was added (Fig. 1; Supplementary Scheme 1). The PEG_24_ linker included an *N*-hydroxysuccinimide ester that allowed reaction with the modified cofactor and a maleimide group for conjugation with the fusion proteins. Conjugation of MAL-PEG_24_-2*AE*-ADP (**11**) and MAL-PEG_24_-2*AE*-NAD^+^ (**13**) to the nanomachines yielded enzymes that were active in batch reactions without the addition of exogenous cofactor, with catalytic constants equivalent or superior to individual enzyme components (Table 1; Supplementary Figure 3). Mass spectrometry of cofactor-conjugated proteolyzed nanomachines indicated that the cysteine in the linker between the catalytic and cofactor recycling modules reacted preferentially with the PEG_24_-maleimide modified cofactors (Supplementary Figure 4). Between 80-100% of the target cysteine residue was conjugated with the modified cofactor for both nanomachines.

### Function of Nanofactory

The nanomachines were immobilized on TFK-activated agarose beads at densities of 1.6, 1.0 and 1.0 milligrams protein per gram wet beads for the phosphorylation, oxidation and aldolase nanomachines, respectively, and packed into glass columns to produce three nanomachine packed bed reactor columns: a phosphorylation column (23.1 mL packed volume), an oxidation column (25.7 mL packed volume) and an aldol addition column (17.7 mL packed volume) (Figure 2).

Individual performance data for each nanomachine reactor (Figure 3) revealed that the maximum space time yields obtained ranged from ∼11 mg L^−1^ h^−1^ mg^−1^ (mg product per mg protein per liter per hour) for the oxidation reactor to ∼70 mg L^−1^ h^−1^ mg^−1^ for the phosphorylation reactor, with the aldol addition reactor yielding ∼30 mg L^−1^ h^−1^ mg^−1^. This is consistent with the expected yields based on *k*_cat_/*K*_M_ values of the loaded enzymes (Table 1), e.g., for the phosphorylation reactor, 36.9 mg protein per column (2.7 nmol tethered biocatalysts) yielded 2.6 g product per litre (6.6 mM) per hour, equivalent to the expected 1,399 nmol per nmol of enzyme per second.

**Figure 3.**
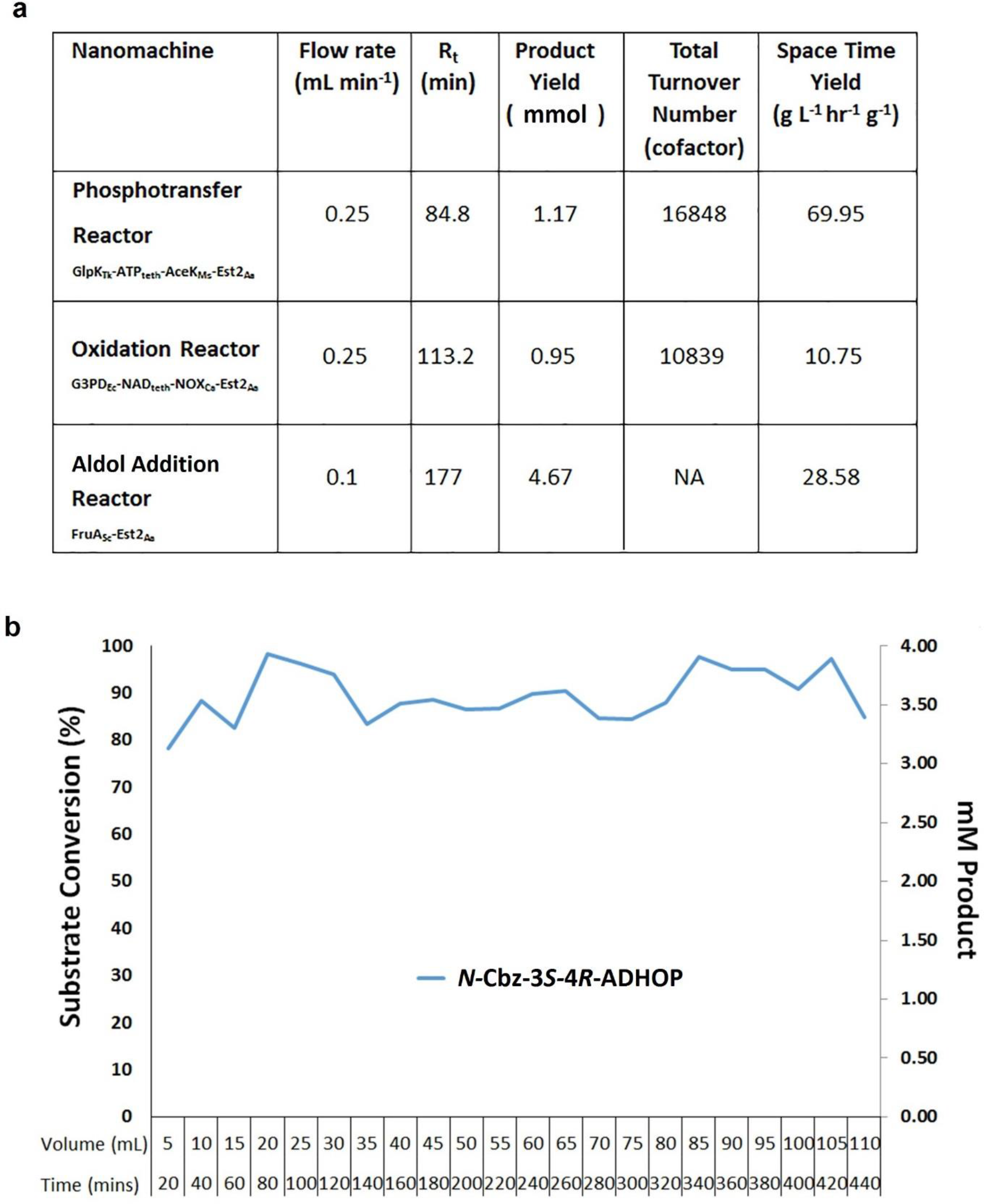
Functional analysis of the three-step nanofactory for the synthesis of D-fagomine from glycerol. **a**, Product yield, cofactor total turnover numbers and space-time yield metrics for each nanomachine reactor. **b**, The nanofactory maintained continuous product yields of between 85-90% conversion of glycerol to *N-Cbz*-3*S*,4*R*-amino-3,4-dihydroxy-2-oxyhexyl phosphate (*N*-Cbz-3*S*,4*R*-ADHOP) for more than 7 h.

As reported elsewhere, when the glycerol-3-phosphate oxidation reaction and aldolase reaction were run in batch, yields were limited by product inhibition and substrate: product equilibrium, with substrate conversion of 88% and 63% respectively ^18, 21, 22^. In our reactors, the phosphorylation and oxidation reactions were run to completion (i.e., complete 100% substrate conversion). It is likely that running the system as a continuous flow reaction prevented the build-up of reaction products and so mitigated both product inhibition and equilibrium control, resulting in the high yields observed in our reactors.

The turnover numbers for the cofactors exceeded 10,000 (∼11,000 for the NAD^+^-dependant oxidation reactor and ∼17,000 for the ATP-dependant phosphorylation reactor; Figure 3a). In each case the reactions stopped because of the inactivation of one of the modules (AceK_Ms_ for the phosphotransfer reactor; NOX_Ca_ for the oxidation reactor) rather than the loss of cofactor. It is reasonable to assume that the turnover numbers would be higher if the enzymes were modified for greater stability, a relatively facile exercise with modern enzyme engineering approaches ^23, 24^.

The three reactors were then combined in series (Figure 2b), with glycerol fed to the first reactor (the phosphorylation reactor) and a feed of Cbz-protected aldehyde entering the system between the oxidation and aldol addition reactors, to yield a three component ‘nanofactory’ for the production of Cbz-protected chiral sugar phosphates from glycerol. When run at 0.3 mL per minute, over 80% of glycerol was converted to enantiomerically pure *N*-Cbz-3*S*,4*R*-ADHOP in a single passage through the reactor (the product was confirmed by HPLC, LCMS and ^1^H NMR analysis; Supplementary Figure 6). The percent conversion dropped to 40% at higher flow rates (1.0 mL.min^−1^), albeit this could be improved through reactor engineering (longer columns, greater biocatalyst loading, multiple passages through the reactor, *etc*.).

## Conclusions

We have developed and successfully implemented a general chemo-genetic protein engineering strategy that enables cofactor-dependent, continuous-flow biocatalysis *via* the use of nanomachines: single molecule multi-enzyme biocatalysts that retain and recycle their cofactors. The engineered biocatalysts were used to construct a three step continuous-flow reactor system (a ‘nanofactory’) that performed well, with superior yields of D-fagomine precursor compared to chemical syntheses ^16^, as well as high space-time yields and total turnover numbers for the catalysts and cofactors. Additionally, use of the biocatalysts in a continuous-flow system appears to have mitigated production inhibition and equilibrium control of yield, allowing very high substrate conversion.

We have used sugar analog synthesis as a model for our prototype ‘nanofactory’; however, we believe that this approach is generalizable because of the modular design principles used in the design of both the ‘nanomachines’ and the ‘nanofactories’. For the nanomachines, we envision a small library of conjugation and cofactor recycling modules that could be used in conjunction with a larger library of catalysis modules to provide access to a wide range of reactions. For example, whilst we have chosen to demonstrate stereoselective aldol addition with the fructose-1,6-biphosphate aldolase FruA, use of the three other classes of DHAP-dependant aldolases (fuculose-1-phosphate FucA, rhamnulose-1-phosphate aldolase RhuA, tagatose-1,6-biphosphate aldolase TagA) ^22^ for the aldol addition reactor could be employed to generate all four ADHOP diastereomers with simple flow-path changes. The molecular modularity of the nanomachines is mirrored in the flow reactor design, providing a flexible platform for building complex, multistep biochemical pathways with both serial and parallel reactor compartments that could be extended into the development of artificial metabolic networks.

## Acknowledgements

We would like to acknowledge the Science and Industry Endowment Fund (SIEF) for funding this work. We would like to thank Drs Matthew Wilding (Australian National University) and John Oakeshott (CSIRO) for their constructive comments during the preparation of this manuscript.

## Author Contributions

CH, CS, CW, NT, JS, GS, GC conceived and designed the study. CH, JS, CW, NF, QI, AN, AF, TN, CN-J performed experiments. CH, CW, JS, AF, AN, TN, QI, CN-J analyzed data and AW performed computational modelling analysis. CH, CW, AN, JS, NF, TN and CS wrote the paper.

## ON-LINE METHODS

### General

Unless otherwise stated in the text, all chemicals were purchased from Sigma-Aldrich (Merck, Australia). For the flow reactors, regulated flow rates and mixing was provided by a modified Biologic DuoFlow system with a Biologic Fraction Collector (Biorad laboratories Inc., USA) for collection of samples. Biologic DuoFlow software v 5.10 Build 2 (Biorad Laboratories Inc., USA) was used to program and control the system, as per manufacturer’s instructions. All restriction enzymes and T4 DNA ligase enzymes used for DNA manipulation were purchased from New England Bioabas (NEB, USA). All PEG compounds were purchased from Quanta BioDesign Ltd (Plain City, OH, USA) and used as received. All other reagents and solvents were obtained from Sigma-Aldrich (Merck), Acros Organics or TCI Chemicals and used as-purchased. Nuclear Magnetic resonance (NMR) spectra were recorded with a Bruker Avance 400 MHz spectrometer in the deuterated solvents as specified. Chemical shifts (δ) were calibrated against residual solvent peaks and are quoted in ppm relative to TMS.

Unless otherwise specified in the text, analytical high-performance liquid chromatography (aHPLC) was performed with a Waters Alliance e2695 Separations Module equipped with Waters 2998 PDA and Acquity QDa detectors, using a Waters XBridge BEH C18 column, 130 Å, 3.5 µm, 2.1 × 50 mm. The following buffer system and gradients were applied: Milli-Q water (Merk Millipore) with 0.1% formic acid (Buffer A) and CH_3_CN with 0.1% formic acid (Buffer B), with a gradient of 0-90% Buffer B unless otherwise specified.

Semi-preparative reversed-phase high performance liquid chromatography (RP-pHPLC) was undertaken using a Shimadzu SPD-10A UV-Vis detector with UV detection at λ 260 nm. The system was equipped with a Shimadzu 322 pump and Gilson 255 Liquid Handler and set at a flow rate of 10 mL min^−1^. Peak separation was achieved using a Vydac 218TP1022 C18, 10 µm, 22 × 250 mm column. Solvent gradients used Milli-Q water with 0.1% TFA (Buffer A) and acetonitrile with 0.1% TFA (Buffer B), with a gradient of 0-70% Buffer B unless otherwise specified.

### DNA manipulation

The pAF1 vector that encodes the DHAP-dependent fructose-1,6-biphosphate aldolase from *Staphylococcus carnosus* was a gift from Dr A. Frazer (University of Manchester, Manchester, UK). For all other constructs, the gene of interest was sourced as described previously ^25^, codon-optimized for expression in *E. coli* and synthesized by GeneArt (ThermoFisher Scientific, Germany), then cloned into pETCC2 ^26^ (a modified version of pET14b, Novagen) using *Nde* I and *Bam* HI or *Eco* RI sites, to create an in-frame *N*-terminal hexa-histidine tag. Genes encoding fusions between the catalytic and cofactor-recycling domains (GlpK_Tk_-AceK_M_s and G3PD_Ec_-NOX_Ca_) were synthesized. Two versions of the synthetic genes were made. In the first version, a single open reading frame containing both domains, separated by a linker and terminating in a STOP codon were synthesized, and included a 5’ *Nde* I site and 3’ *Bam* HI site for cloning. The second version differed in that the stop codon was omitted and a 3’ *Sph* 1 site was included to allow construction of the fusions that included the conjugation domain. The versions with a STOP codon were cloned into pETCC2 for expression and purification of GlpK_Tk_-AceK_M_s and G3PD_Ec_-NOX_Ca_, whilst the *Nde* I-*Sph* I versions were subcloned into a prepared pETCC2 backbone containing the esterase *e*2_Aa_ gene to create a genetic fusion via the *Sph* I site, with a short Gly-Ser repeat spacer. This yielded genes encoding the GlpK_Tk_-AceK_M_s-E2_Aa_ and G3PD_Ec_-NOX_Ca_-Est2_Aa_ nanomachines. A similar strategy was used to fuse *fru*A_Sc_ with *e*2_Aa_ via the *Sph* I site yielding a construct encoding FruA_Sc_-Est2_Aa_ (Supplementary Figure 1). The final insertion fragments used for each of the nanomachines are depicted in Supplementary Figure 1. All constructs were confirmed by DNA sequencing (Macrogen, S. Korea).

**Supplementary Figure 1.**
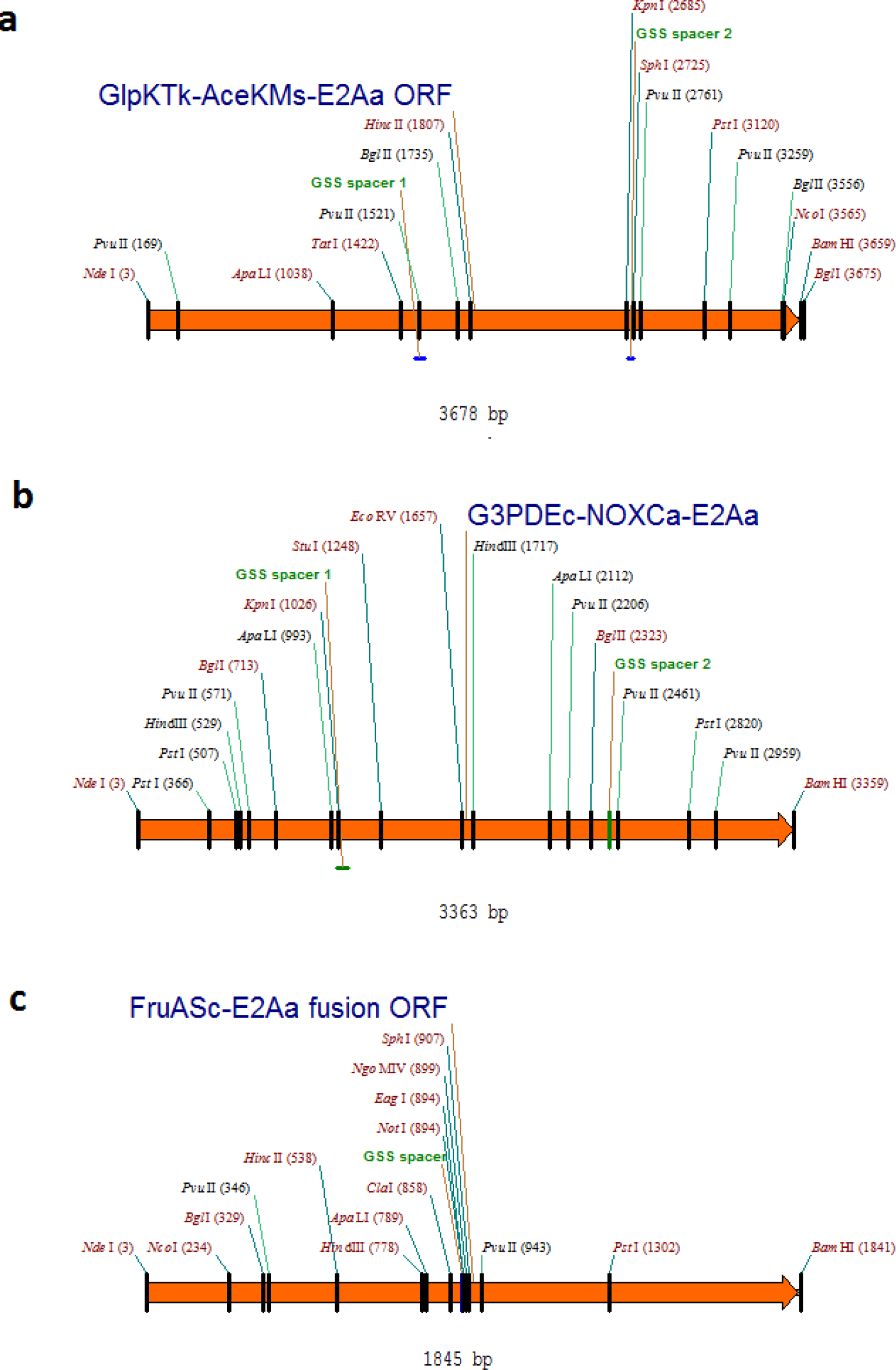
**DNA insertion regions for each of the nanomachine expression constructs** encoding GlpK_Tk_-AceK_M_s-E2_Aa_ (a), G3PD_Ec_-NOX_Ca_-Est2_Aa_ (b) and FruA_Sc_-Est2_Aa_ (c), combining each multienzyme fusion as a single ORF for insertion into pETCC2 via the *Nde* 1-*Bam* H1 sites to create an in-frame 5’-hexahistidine tag.

### Engineered Fusion Protein Sequences

The hexaHIS-tagged individual and fusion protein sequences used in this study are listed below, with linker regions between the protein domains highlighted in bold and underlined, and the cysteine required for tethering modified cofactor highlighted in red bold font:

#### G3PD_Ec_

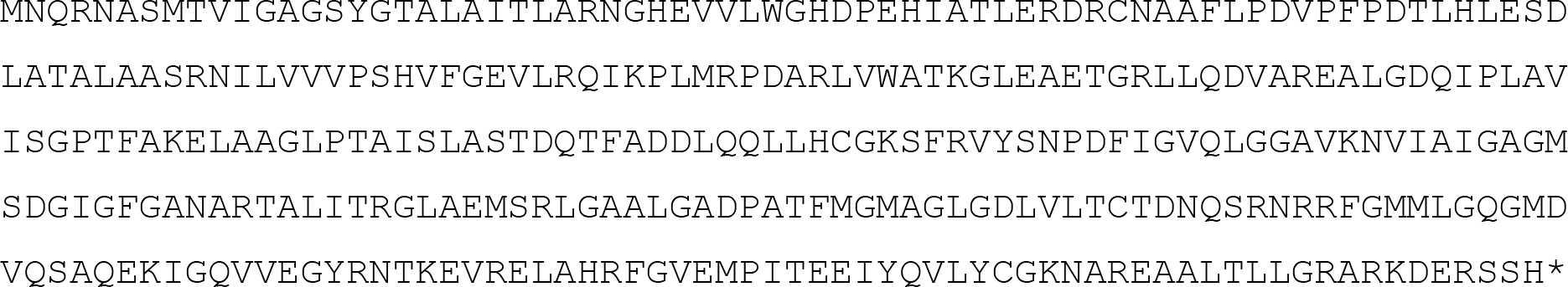

#### Nox_Ca_

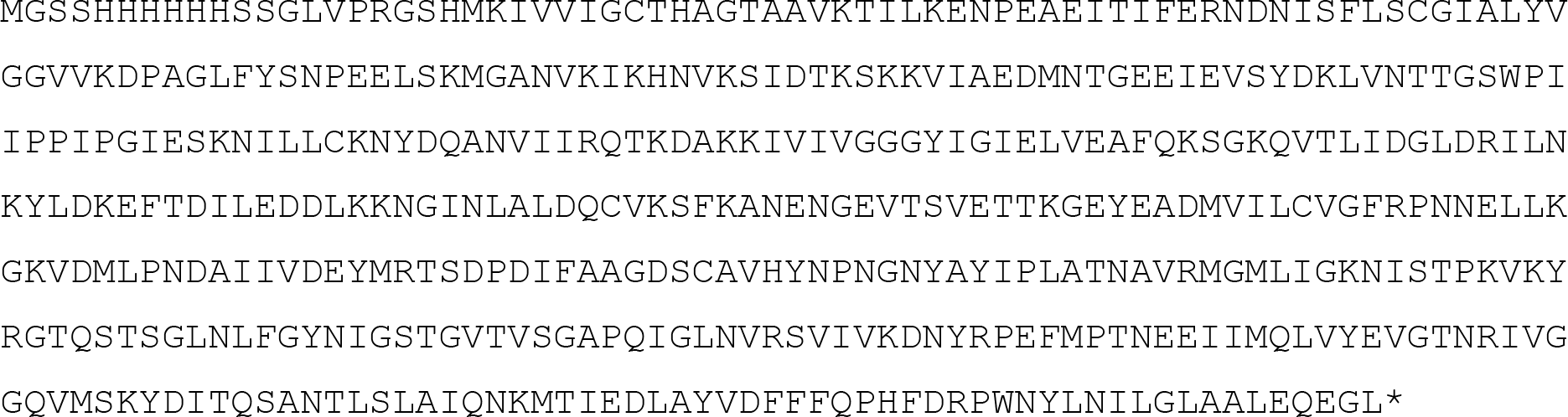

#### GlpK_Tk_

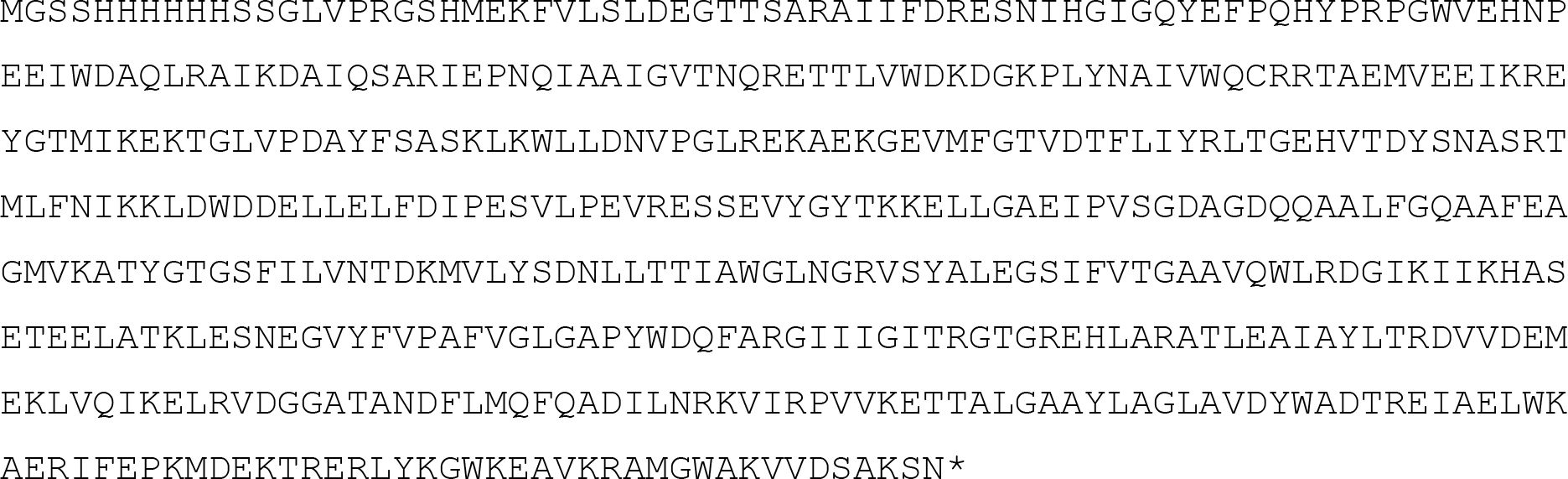

#### AceK_Ms_

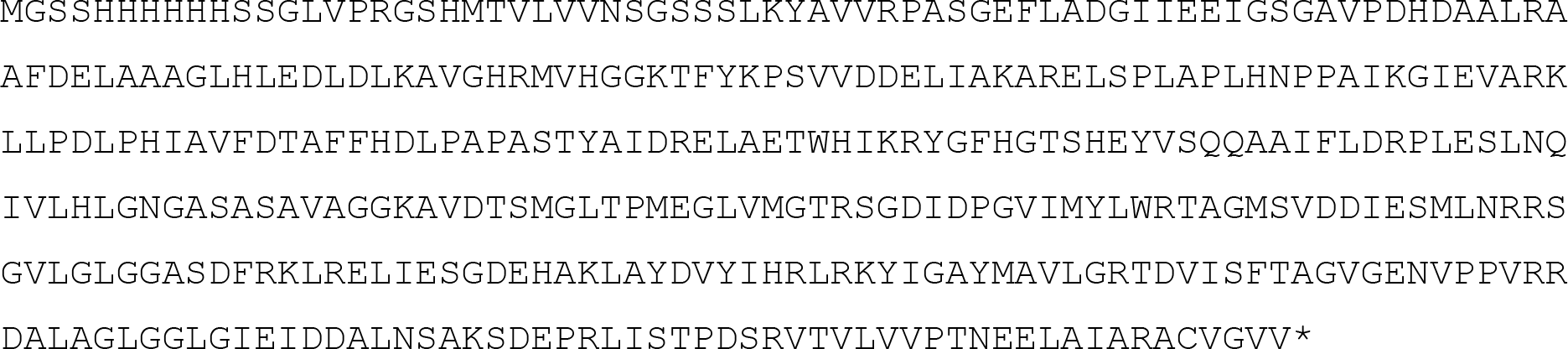

#### GlpK_Tk_-AceK_M_s

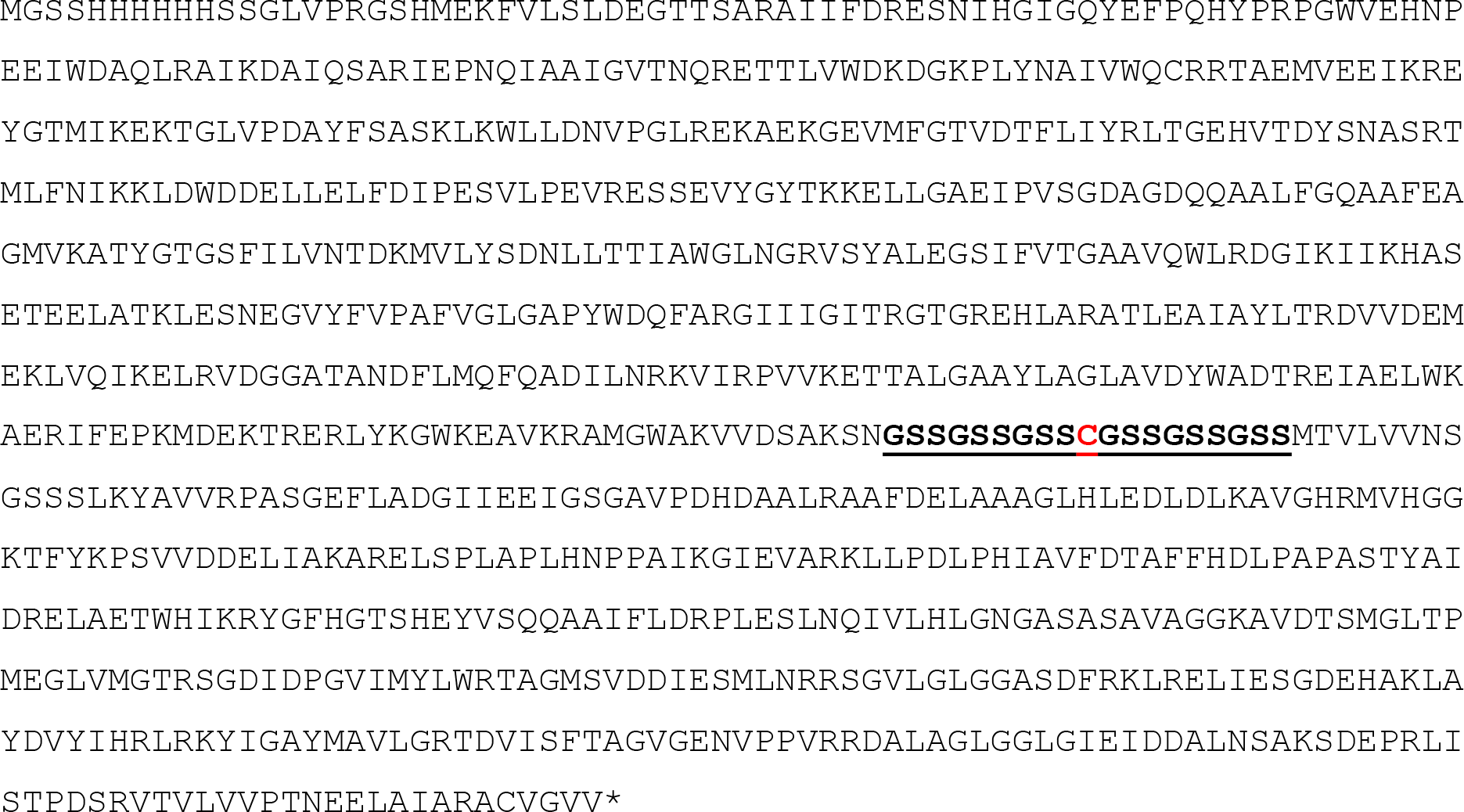

#### G3PD_Ec_-NOX_Ca_

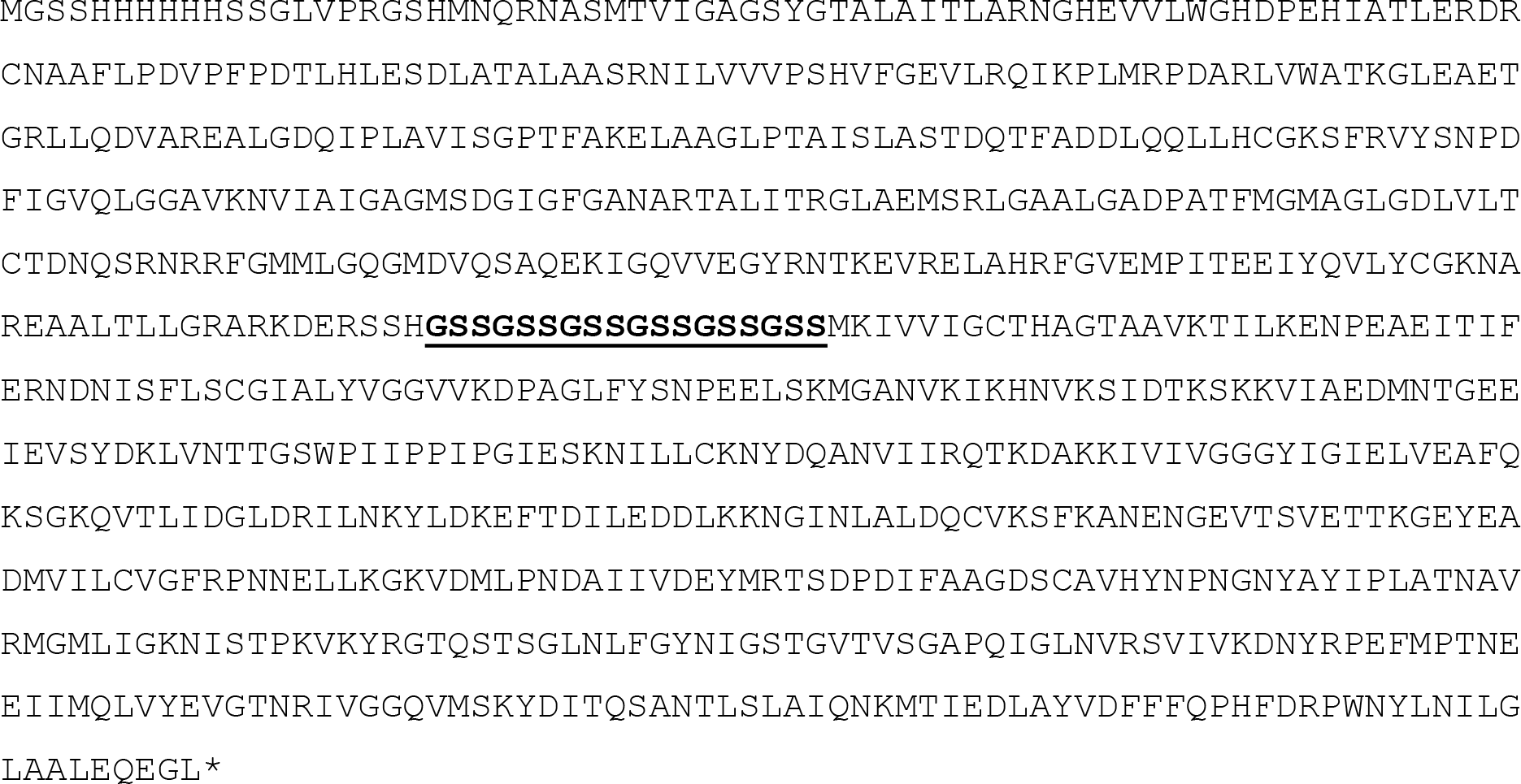

#### GlpK_Tk_-AceK_M_sE2_Aa_

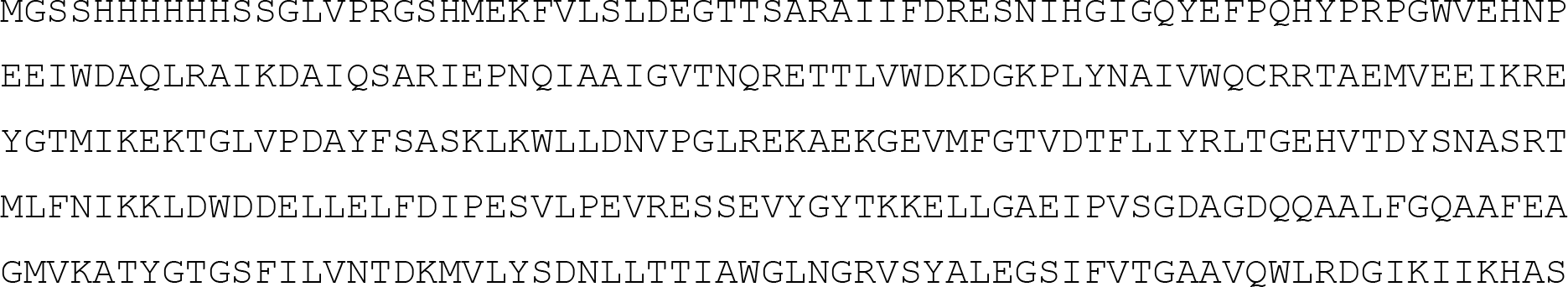

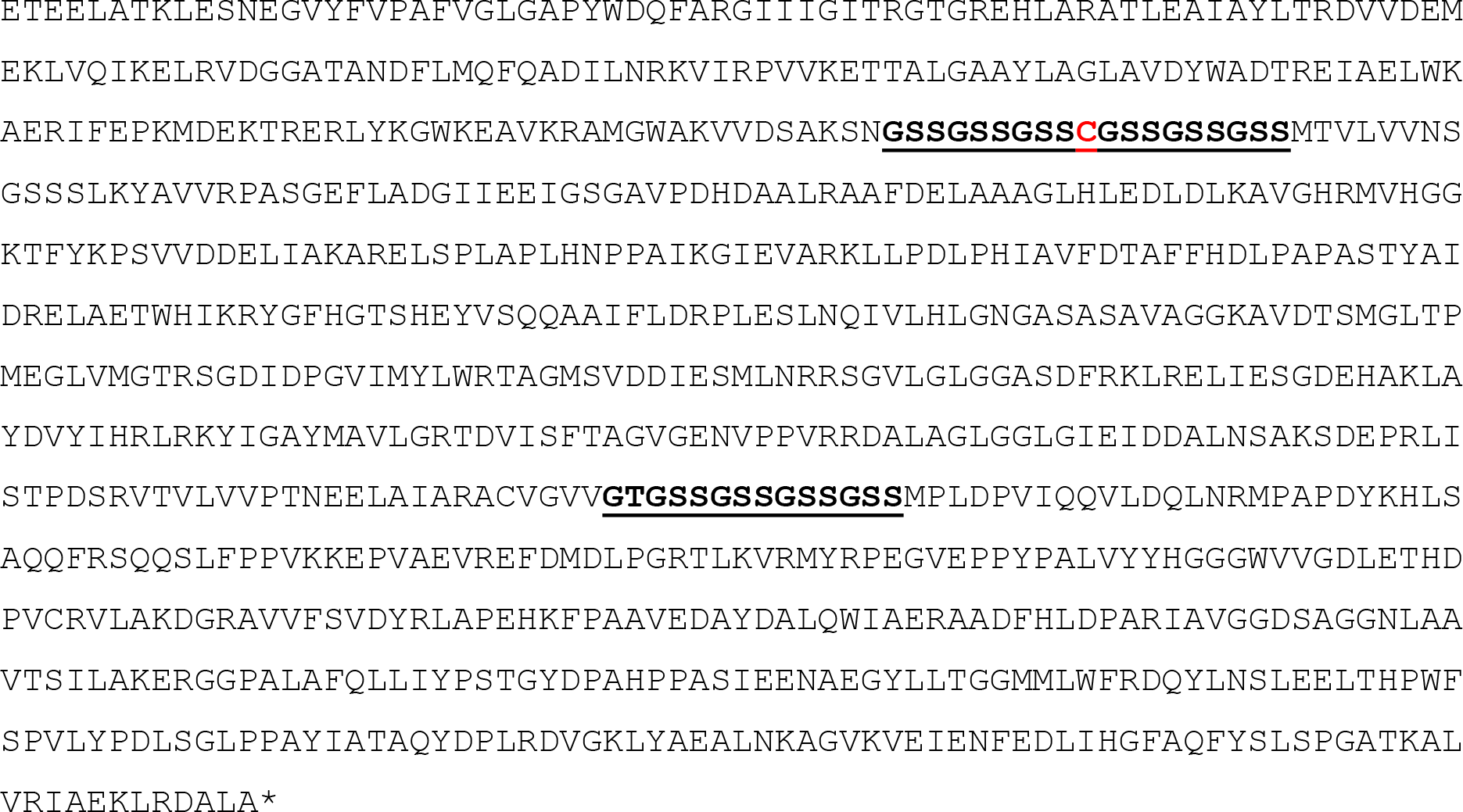

#### G3PD_Ec_-NOX_Ca_-Est2_Aa_

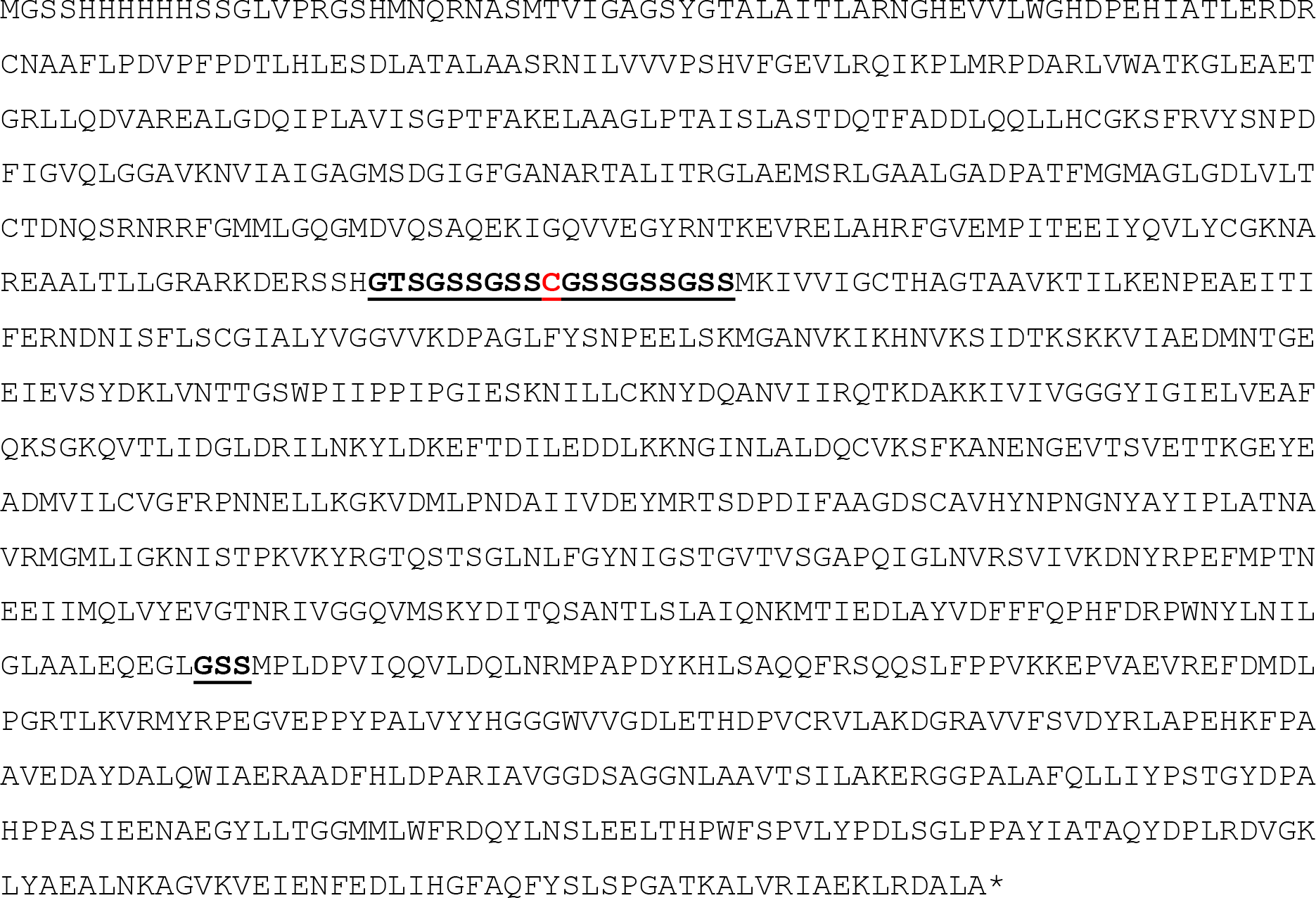

#### FruA_Sc_-Est2_Aa_

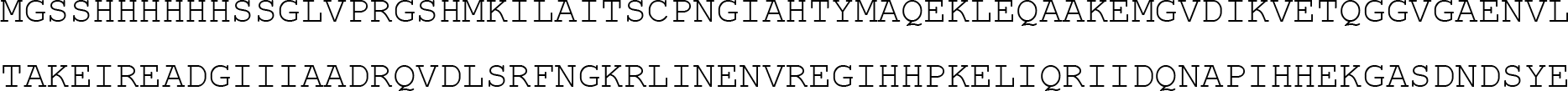

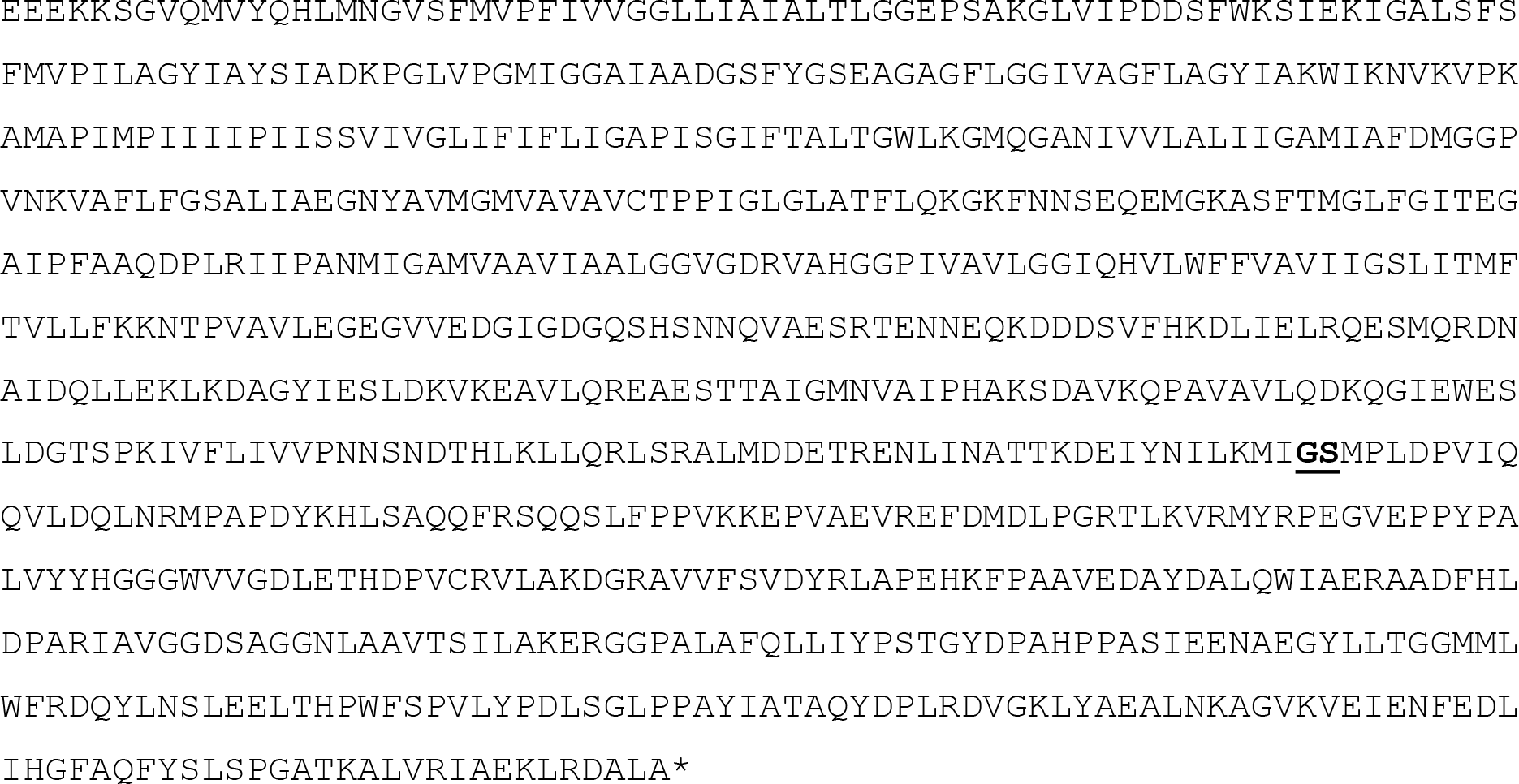

### Protein Expression and Purification

#### Expression of individual enzymes and bi-enzymatic fusion proteins

The expression plasmids outlined above were used to transform *E. coli* BL21 DE3 Star (Invitrogen, ThermoFisher Scientific, USA), using Luria agar containing 100 µg mL^−1^ ampicillin as a selective growth medium. Cells were cultured overnight in Luria broth containing 100 µg mL^−1^ ampicillin at 37 °C and shaken at 200 rpm, then induced for 2, 4, 6 and 24 h with either arabinose or isopropyl β-D-1-thiogalactopyranoside (IPTG) at 0.2 M and 1 mM final concentration, respectively (see Supplementary Table 1 for details). Cultures were then harvested, by centrifugation at 8000 g, resuspended in one tenth culture volume of resuspension buffer (50 mM Tris-Cl, 250 mM NaCl, pH 7.5) and lyzed with Bugbuster™ (Novagen). Protein expression was analyzed by SDS-PAGE separation (4-12% Bolt Bis-Tris Plus Polyacrylamide Gel with MES SDS running buffer (Invitrogen, USA) and visualized with NuBlue (Novagen). The optimal expression time (Supplementary Table 1) was selected and large scale expression cultures of 1-2 L prepared in the same way as above except that cells were lysed by passage through an EmulsiFlex-C5 cell homogenizer (Avestin) at 20,000 psi, 4 °C and cellular debris removed by centrifugation (40,000 × *g*, 15 min, 4 °C). Protein was first purified from cell free lysates by IMAC purification of HIS-tagged protein by elution with resuspension buffer (50 mM Tris-Cl, 250 mM NaCl, pH 7.5) containing increasing concentration of imidazole from NiNTA-sepharose (Hi5 HIS-TRAP, GE Healthcare). The desired protein fractions were then pooled and further purified using a Superdex 200 size exclusion column (GE Healthcare). Pooled fractions were then concentrated and stored at 4 °C, or − 80 °C, as required.

**Supplementary Table 1.** Optimal recombinant expression conditions for the individual enzymes and multi-enzyme fusions comprising each nanomachine.

| Enzyme | Construct | Expression Vector | Host Cells | Inducer; Temperature (°C) and Time [h] |
| --- | --- | --- | --- | --- |
| **Phosphotransfer Nanomachine Proteins** |  |  |  |  |
| GlpK<sub>TK</sub> | pCJH1 | pETCC2 | *E.coli* BL21DE3 Star (Invitrogen) | 1mM IPTG; 37°C, 18h |
| AceK<sub>MS</sub> | pCJH2 | pDEST17 (Invitrogen) | *E.coli* BL21AI (Invitrogen) | 20mM arabinose 15°C, 18h |
| GlpK<sub>TK</sub>-AceK<sub>MS</sub> | pCJH3 | pETCC2 | *E.coli* BL21DE3 Star (Invitrogen) | 1mM IPTG; 37°C, 18h |
| GlpK<sub>TK</sub>-AceK<sub>MS</sub>-Est2<sub>Aa</sub> | pCJH4 | pETCC2 | *E.coli* BL21DE3 Star (Invitrogen) | 1mM IPTG; 37°C, 18h |
| **Oxidation Nanomachine Proteins** |  |  |  |  |
| G3PD<sub>Ec</sub> | pCJH5 | pDEST17 (Invitrogen) | *E.coli* BL21AI (Invitrogen) | 20mM arabinose 25°C, 18h |
| NOX<sub>Ca</sub> | pCJH6 | pETCC2 | *E.coli* BL21DE3 Star (Invitrogen) | 1mM IPTG; 37°C, 18h |
| G3PD<sub>Ec</sub>-NOX<sub>Ca</sub> | pCJH7 | pETCC2 | *E.coli* BL21DE3 Star (Invitrogen) | 1mM IPTG; 25°C, 18h |
| G3PD<sub>Ec</sub>-NOX<sub>Ca</sub>-Est2<sub>Aa</sub> | pCJH8 | pETCC2 | *E.coli* BL21DE3 Star (Invitrogen) | 1mM IPTG; 25°C, 18h |
| **Aldol addition Nanomachine Proteins** |  |  |  |  |
| FruA<sub>Sc</sub> | pAF1 | pRSET-A (Invitrogen) | *E.coli* BL21DE3 | 1mM IPTG; 15°C, 18h |
| FruA<sub>Sc</sub>-Est2<sub>Aa</sub> | pCJH9 | pETCC2 | *E.coli* BL21DE3 Star (Invitrogen) | 1mM IPTG; 15°C, 18h |
| **Esterase Conjugation Domain Protein** |  |  |  |  |
| Est2<sub>Aa</sub> | pCJH10 | pETCC2 | *E.coli* BL21DE3 Star (Invitrogen) | 1mM IPTG; 37°C, 18h |

Each of the purified individual and bi-enzymatic fusion proteins were then characterised in terms of catalytic activity (Table 1), thermostability and oligomeric structure (Supplementary Table 2) to ensure suitability for incorporation into the final nanomachine constructs with the esterase conjugation domain, utilising the enzymatic activity assays and analytical methods outlined below. Further specific details regarding the expression and purification of the final three nanomachine multi-enzyme fusion proteins used to construct the nanofactory is given below.

**Supplementary Table 2.** Biochemical characterization of the individual enzymes and multi-enzyme fusions comprising each nanomachine.

| | Optimal<br>Reaction pH<br>(pH range) | $T_{50}^*$<br>(°C) | Oligomeric Structure | $k_{cat}/K_M$<br>($s^{-1} M^{-1}$) | Reference |
| --- | --- | --- | --- | --- | --- |
| **Glycerol phosphorylation** |  |  |  |  |  |
| GlpK<sub>TK</sub> | 8.0 (6.5-9.5) | > 100 | monomer | $6.1 \times 10^7$ | <sup>27</sup>; This study |
| GlpK<sub>TK</sub>-AceK<sub>MS</sub> | | 58 | dimer | $7.7 \times 10^7$ | This study |
| GlpK<sub>TK</sub>-AceK<sub>MS</sub>-Est2<sub>Aa</sub> | | 59 | monomer/hexamer | $8.6 \times 10^7$ | This study |
| **Acetate dephosphorylation (ADP phosphorylation)** |  |  |  |  |  |
| AceK<sub>MS</sub> | 7.4 (6.0-8.5) | 52 | homodimer | $2.8 \times 10^6$ | <sup>28</sup>; This study |
| GlpK<sub>TK</sub>-AceK<sub>MS</sub> | | 50 | dimer | $5.4 \times 10^5$ | This study |
| GlpK<sub>TK</sub>-AceK<sub>MS</sub>-Est2<sub>Aa</sub> | | 63 | monomer/hexamer | $9.1 \times 10^5$ | This study |
| **Glycerol-3-phosphate oxidation** |  |  |  |  |  |
| G3PD<sub>EC</sub> | 9.0 (8.0-9.5) | 51 | monomer | $1.4 \times 10^6$ | <sup>18</sup>; This study |
| G3PD<sub>EC</sub>-NOX<sub>Ca</sub> | | 37 | dimer | $1.8 \times 10^4$ | This study |
| G3PD<sub>EC</sub>-NOX<sub>Ca</sub>-Est2<sub>Aa</sub> | | 45 | dimer | $1.1 \times 10^4$ | This study |
| **Oxygen reduction (NADH oxidation)** |  |  |  |  |  |
| NOX<sub>Ca</sub> | 7.0 (7.0-9.0) | 37 | homodimer | $4.9 \times 10^6$ | <sup>29</sup>; This study |
| G3PD<sub>EC</sub>-NOX<sub>Ca</sub> | | 37 | dimer | $6.2 \times 10^6$ | This study |
| G3PD<sub>EC</sub>-NOX<sub>Ca</sub>-Est2<sub>Aa</sub> | | 37 | dimer | $4.6 \times 10^6$ | This study |
| **Aldol addition** |  |  |  |  |  |
| FruA<sub>Sc</sub> | 6.5-9.0 | > 95 | monomer | $3.2 \times 10^4$ | <sup>30</sup> |
| FruA<sub>Sc</sub>-Est2<sub>Aa</sub> | | 81 | monomer | $1.3 \times 10^5$ | This study |
| **Esterase<sup>1</sup>** |  |  |  |  |  |
| Est2<sub>Aa</sub> | 7.1 (5.5-8.0) | 80 | monomer | $8.3 \times 10^5$ | <sup>31</sup>; This study |

\* $T_{50}$ the temperature at which, after 30 min of incubation, 50% of the initial enzyme activity remains. The $T_{50}$ values are the averages of at least three independent assays. Standard deviations were below 1.0 °C in all cases. pH optima did not vary between individual and fusion enzymes.

#### Expression of GlpK_Tk_-AceK_Ms_-E2_Aa_

The expression of GlpK_*Tk*_-AceK_*M*_s-E2_*Aa*_ in *E. coli* BL21 DE3 Star (Invitrogen, ThermoFisher Scientific, USA) cells transformed with the plasmid pETCC2-GlpK_*Tk*_-AceK_*M*_s-E2_*Aa*_ (pCJH4; Supplementary Table 1) was examined after induction at 15 °C for 18 h with 1 mM IPTG as inducer, and the oligomeric state related to fusion protein activity, using glycerol kinase activity as a proxy for all three activities (Supplementary Figure 2). Although all three oligomeric states isolated by gel filtration purification (Superdex 150 gel filtration column, GE Healthcare, after pooling HIS-tagged purification pools from a 5 mL HisTrap FF column, GE Healthcare) demonstrated glycerol kinase activity, the maximum specific activity per milligram of protein was retained in the “monomeric-dimeric” fraction (Supplementary Figure 2). For subsequent experiments, the monomeric-dimeric fraction was isolated.

For large scale preparation of GlpK_*Tk*_-AceK_*M*_s-E2_*Aa*_ fusion protein pCH4 was transformed into *E. coli* BL21DE3 Star cells. Cells were cultured in Luria broth overnight at 37 °C with shaking at 200 rpm, diluted to OD_600nm_ 0.7 in Luria broth and induced at 15 °C for 18 h with arabinose and IPTG (20 mM and 1 mM final concentration, respectively) and then harvested, washed in one tenth volume resuspension buffer (50 mM Tris-Cl, 250 mM NaCl, pH 7.5) and cell pellets stored at −20 °C. Cell paste (8 g) was resuspended in 200 mL 50 mM Tris, 300 mM NaCl pH 8 containing 0.5 mg mL^−1^ lysozyme (Sigma–Aldrich), 2 mM PMSF (Sigma–Aldrich), four EDTA-Free Complete Protease inhibitor tablets (Roche) and 1000 Units Benzonase (Merck Millipore). Following resuspension, the cells were ruptured by passage three times through an EmulsiFlex-C5 cell homogenizer (Avestin) at 15,000 psi at 4 °C and cellular debris removed by centrifugation (40,000 × *g*, 15 min, 4 °C). The lysate was filtered (0.45 μm) and one quarter applied to a 5 mL HisTrap FF column (GE Healthcare) equilibrated in 50 mM Tris, 300 mM NaCl pH 8 containing 0.1 mM tris-(2-carboxyethyl)phosphine (TCEP). The column was washed with 40 mM imidazole in the same buffer then the bound protein eluted with 300 mM imidazole in the same buffer. The eluted protein was analyzed by gel filtration on a Superdex 200 1030 gel filtration column (GE Healthcare) equilibrated with phosphate buffered saline with the absorbance of the eluted protein monitored at 280 nm and the esterase activity in the eluted fractions determined. For comparison, 0.5 mL of crude lysate was also subjected to gel filtration analysis, with monitoring at 280 nm and analysis of esterase activity in the fractions.

#### Expression and purification of G3PD_Ec_-NOX_Ca_-Est2_Aa_

Briefly, *E.coli* BL21 DE3 Star (Invitrogen) cells expressing G3PD_Ec_-NOX_Ca_-Est2_Aa_ (pCJH8, Supplementary Table 1) were cultured in an XRS 20 bioreactor (Pall Corporation, USA) using a 2 litre volume of M9 minimal medium, with 1% (w/v) ammonium sulphate and 1% (w/v) glucose as nitrogen and carbon source respectively and supplemented with 100 µg mL^−1^ ampicillin. After initial growth at 37 °C, the temperature was reduced to 25 °C prior to induction, when the OD_600nm_ reached 2.2. The optical density of the culture at induction was OD_600nm_ 2.9 and 1.6 mL of 20% arabinose and IPTG to 1 mM were added to induce. The glucose feed was started 7 h post-induction to maintain 1% glucose and cells were harvested 22 h post-induction, when OD_600nm_ was 21.6.

Cell paste (2 g) was resuspended in 50 mL 50 mM Tris, 300 mM NaCl pH 8 containing 0.1 mM Tris-(2-carboxyethyl)phosphine (TCEP; Sigma-Aldrich), 0.5 mg mL^−1^ lysozyme (Sigma–Aldrich), 2 mM phenylmethane sulfonyl fluoride (PMSF; Sigma–Aldrich), one EDTA-Free Complete Protease inhibitor tablet (Roche) and 250 Units Benzonase (Merck Millipore). Following resuspension, the cells were ruptured by passage three times through an EmulsiFlex-C5 cell homogenizer (Avestin) at 15,000 psi at 4 °C and cellular debris removed by centrifugation (40,000 × *g*, 15 min, 4 °C). The lysate was filtered (0.45 μm) and applied to a 5 mL HisTrap FF column (GE Healthcare) equilibrated in 50 mM Tris, 300 mM NaCl pH 8 containing 0.1 mM TCEP. The column was washed with 40 mM imidazole in the same buffer then the bound protein eluted with 300 mM imidazole in the same buffer. The eluted protein was subjected to gel filtration on a Superdex 200 gel filtration column (GE Healthcare) equilibrated with 50 mM citrate, 200 mM NaCl pH 6 containing 1 mM TCEP with the absorbance of the eluted protein monitored at 280 and 450 nm. Fractions eluting from 158 – 192 mL were pooled and concentrated to 0.94 mg mL^−1^.

G3PD_Ec_-NOX_Ca_-Est2_Aa_ eluted in a broad peak from the gel filtration column, with some protein eluting in the void volume (Supplementary Figure 2). The final pool was > 95% pure as estimated by SDS-PAGE and was found to have specific activities of 16 U mg^−1^ for esterase and 31 U mg^−1^ for NADH oxidase.

#### Expression and purification of FruA_Sc_-Est2_Aa_

FruA_*Sc*_ was selected as the preferred aldolase for the nanofactory based on its enantioselectivity (3*S*, 4*R*-ADHOP), oligomeric structure, stability, catalytic rate (Supplementary Table 2) and previous success using this enzyme in multi-enzyme cascades to produce similar chiral sugars ^18^.

For the preparation of the aldol addition nanomachine, we constructed genetic fusions between Fru_Sc_ and E2_Aa_ as illustrated in Supplementary Figure 1. Purified Fru_Sc_-E2_Aa_ was obtained by expression and purification from *E. coli* BL21 DE3 Star cells (Invitrogen, Thermofisher Scientific). Briefly, a synthetic gene encoding FruA_Sc_-E2_Aa_ was transferred into pETCC2 (pCJH10, Supplementary Table 1) and used to transform *E.coli* BL21DE3 Star (Invitrogen) cells. Cells were cultured in Luria broth overnight at 37 °C with shaking at 200 rpm, diluted to OD_600nm_ 0.7 in Luria broth and induced for 18 h with arabinose and IPTG (20 mM and 1 mM final concentration, respectively). The cells were then harvested, washed in one tenth volume resuspension buffer (50 mM Tris-Cl, 250 mM NaCl, pH 7.5) and cell pellets stored at −20 °C. Cell paste (8 g) was resuspended in 200 mL 50 mM Tris, 300 mM NaCl pH 8 containing, 0.5 mg mL^−1^ lysozyme (Sigma–Aldrich), 2 mM PMSF (Sigma–Aldrich), four EDTA-Free Complete Protease Inhibitor tablets (Roche) and 1000 Units Benzonase (Merck Millipore). Following resuspension, the cells were ruptured by passage three times through an EmulsiFlex-C5 cell homogenizer (Avestin) at 15,000 psi and 4 °C and cellular debris removed by centrifugation (40,000 × *g*, 15 min, 4 °C). The lysate was filtered (0.45 μm) and applied to a 5 mL HisTrap FF column (GE Healthcare) equilibrated in 50 mM Tris, 300 mM NaCl pH 8 containing 0.1 mM TCEP. The column was washed with 40 mM imidazole in the same buffer then the bound protein eluted with 300 mM imidazole in the same buffer. The eluted protein was analyzed by gel filtration on a Superdex 200/1030 gel filtration column (GE Healthcare) equilibrated with phosphate buffered saline with the absorbance of the eluted protein monitored at 280 nm (Supplementary Figure 2) and the esterase activity in the eluted fractions determined.

**Supplementary Figure 2.**
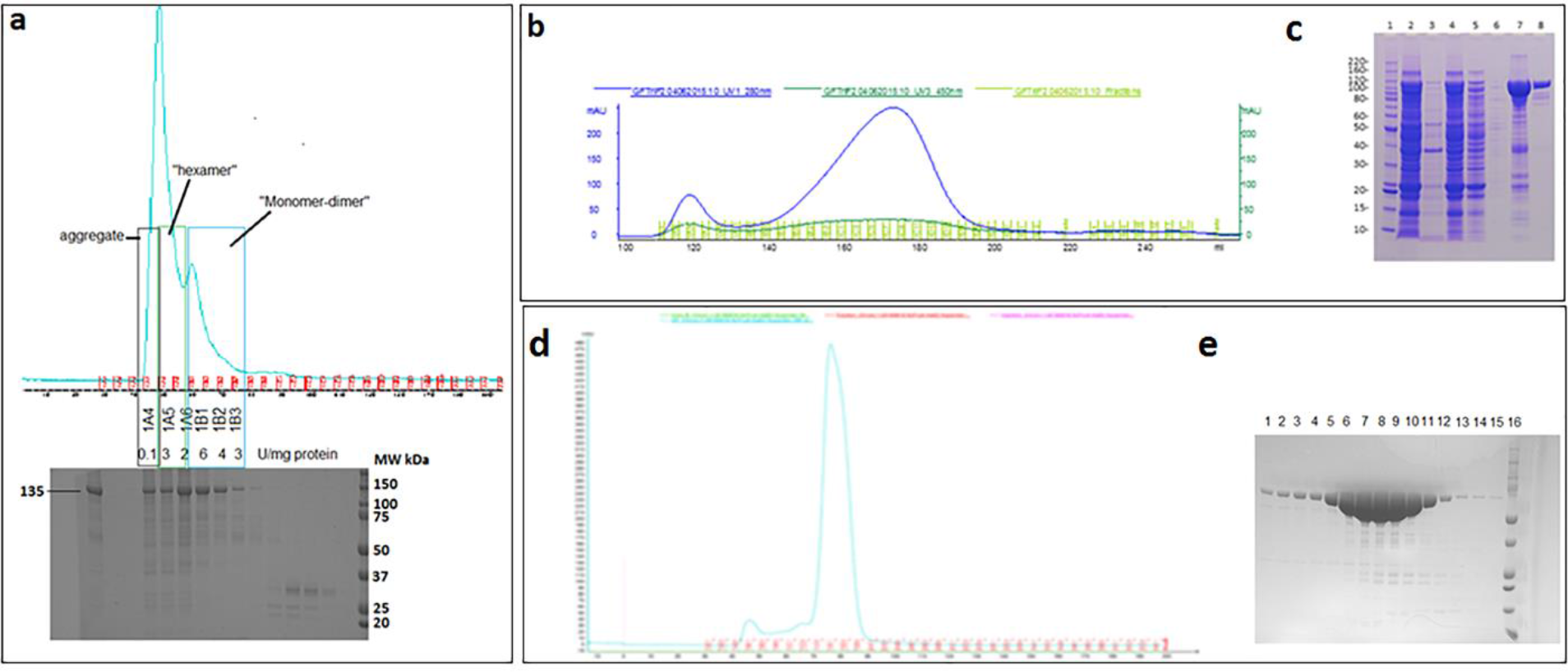
Expression and purification of nanomachine fusion proteins. **a,** Size exclusion fractionation of HIS-tag purified recombinant GlpK_TK_-AceK_Ms_-E2_Aa_ multi-enzyme fusion protein was used to estimate oligomeric state and associated specific activity after induction with 1 mM IPTG at 15 °C for 24 h. **b,** Size exclusion fractionation of HIS-tag purified recombinant G3PD_Ec_-NOX_Ca_-Est2_Aa_. Fractions from 158 – 192 mL were pooled to avoid higher MW aggregate. **c,** SDS-PAGE analysis of G3PD_Ec_-NOX_Ca_-Est2_Aa_ purification. Lane 1 Benchmark™ protein molecular weight standard standard (Invitrogen), Lane 2 whole cells, Lane 3 pellet after centrifugation of lysate, Lane 4 lysate, Lane 5 HisTrap unbound, Lane 6 40 mM imidazole wash, Lane 7 300 mM imidazole elution, Lane 8 gel filtration pool. **d,** Size exclusion fractionation profile of HIS-tag purified recombinant FruA_Sc_-Est2_Aa._ **e,** SDS-PAGE analysis of FruA_Sc_-Est2_Aa_ purification. Lane 1 Fraction 1B1, Lanes 2-15 fractions 1C1 to 2A3, Lane 16 Dual™ Protein molecular weight standard (NEB).

### Cofactor modification

#### Synthesis of *N*^6^-2*AE*-ADP

Adenosine-5’-phosphate sodium salt, ADP.xNa, (0.5 g, 1.13 mmol) was dissolved in distilled deionised (DI) water (1.5 mL) and ethyleneimine (160 µL) was added very slowly. During the addition of the ethyleneimine, the pH was carefully adjusted with perchloric acid to keep it within the pH range of 2.0-4.0. The pH at the end was left at 3.2. The reaction was stirred at room temperature for ∼50 h. The solvent was then evaporated in a fume-hood under a stream of nitrogen. The crude residue was dissolved in DI water (10 mL) and the pH adjusted to 5.6 by the addition of lithium hydroxide (LiOH; saturated aq. solution) and heated at 35 °C for 80 h. The solution was lyophilized to yield crude *N*^6^-2*AE*-ADP (Supplementary Scheme 1).

#### Synthesis of MAL-PEG_24_-2*AE*-ADP

Crude *N*^6^-2*AE*-ADP DIPEA salt (100 mg, crude estimate equivalent to about 13.7 mmol pure 2*AE*-ADP) was dissolved in 50% acetonitrile/phosphate buffered saline pH 7.0 (3.0 mL) and a solution of MAL-PEG_24_-NHS (43.0 mg) was added and stirred at room temperature overnight (Supplementary Scheme 1). The mixture was purified by pHPLC and lyophilised to yield pure MAL-PEG_24_-2*AE*-ADP (12.2 mg).

#### Synthesis of*N*^6^-2*AE*-NAD^+^

To a solution of β-nicotinamide adenine dinucleotide hydrate, NAD^+^, (1.0 g, 1.51 mmol) dissolved in 2 mL deionized water was added dropwise ethyleneimine (4.25 mmol) with the solution maintained at a pH of 3.2 with the addition of 70% perchloric acid. The reaction mixture was stirred at room temperature for 50 h with the pH maintained from 2-3, before the addition of 1.75 mL deionized water to solubilise precipitate. The product was precipitated by the addition of ice-cold ethanol and the precipitate washed with ethanol. The resulting mix of *N*1-2*AE*-NAD^+^ and NAD^+^ was dissolved in water (10 mL) and adjusted to pH 6.5 with 0.1 M LiOH. The solution was stirred at 50 °C for 7 h with the pH maintained at 6.5 before being lyophilized to yield the product, as a mixture of *N^6^*-2*AE*-NAD^+^ and NAD^+^ (Supplementary Scheme 1).

#### Synthesis of MAL-PEG_24_-2*AE*-NAD^+^

To a stirred solution of *N*^6^-2*AE*-NAD^+^/NAD^+^ (14.7 mg mix, approximately 0.0104 mmol *N*^6^-2*AE*-NAD^+^) in phosphate buffered saline (PBS; pH 7.4, 1.0 mL) was added a solution of MAL-PEG_24_-NHS (17.4 mg, 0.0124 mmol) in PBS (1 mL). The solution was stirred at room temperature overnight (Supplementary Scheme 1). The mixture was analyzed by HPLC (0→50% CH_3_CN + 0.1% TFA over 18 min). Rt 17.8 min ESI+ found 662.62 (M/3, calcd 662.65) and 993.42 (M/2, calcd 993.98). The mixture was purified by preparative HPLC and fractions at Rt 17.8 min combined and lyophilized to yield pure MAL-PEG_24_-2*AE*-NAD^+^ (5.4 mg, 26%).

**Supplementary Scheme 1.**
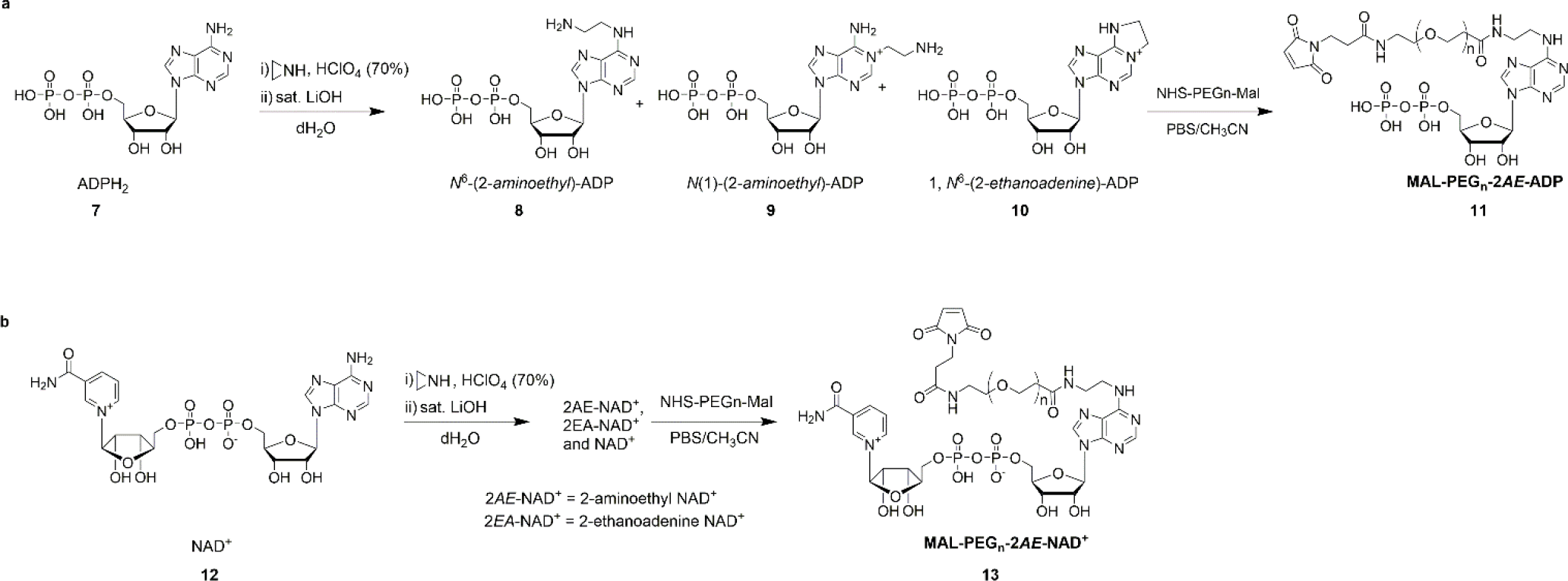
Synthesis of modified cofactors for tethering to nanomachine fusion proteins. **a**, Scheme for synthesis of MAL-PEG_n_-2*AE*-ADP (**11**) from ADP (**7**). **b**, Scheme for synthetic route to prepare MAL-PEG_n_-2*AE*-NAD^+^ (**13**) from NAD^+^ (**12**).

### Cofactor attachment

#### Tethering of MAL-PEG_24_-2*AE*-ADP to GlpK_TK_-AceK_Ms_-Est2_Aa_ in solution

IMAC purified GlpK_TK_-AceK_Ms_-Est2_Aa_ was found to elute as high molecular weight protein and in PBS (Supplementary Figure 2). The IMAC purified protein was treated with TCEP (0.1 mM) then reacted with 10 equivalents of MAL-PEG_24_-2*AE*-ADP without removal of the TCEP and washed with PBS. The tethered 2*AE*-ADP-PEG_24_-MAL-GlpK_TK_-AceK_Ms_-Est2_Aa_ was found to convert 10 mM glycerol and 10 mM acetyl phosphate to glycerol-3-phosphate with high efficiency (Supplementary Figure 3a).

#### Tethering of MAL-PEG_24_-2*AE*-NAD^+^ to G3PD_Ec_-NOX_Ca_-Est2_Aa_ in solution

G3PD_Ec_-NOX_Ca_-Est2_Aa_ was desalted into PBS containing 0 mM, 0.1 mM or 1 mM TCEP and reacted with 1–200 equivalents of MAL-PEG24-2*AE*-NAD^+^. The reaction mixtures were analyzed by SDS-PAGE and the conjugate from one condition (0.1 mM TCEP, 200 equivalents TCEP) analyzed by mass spectrometry. The tethered 2*AE*-NAD^+^-PEG_24_-MAL-G3PD_Ec_-NOX_Ca_-Est2_Aa_ was found to convert 10 mM glycerol-3-phosphate and to DHAP with high efficiency (Supplementary Figure 3b).

**Supplementary Figure 3.**
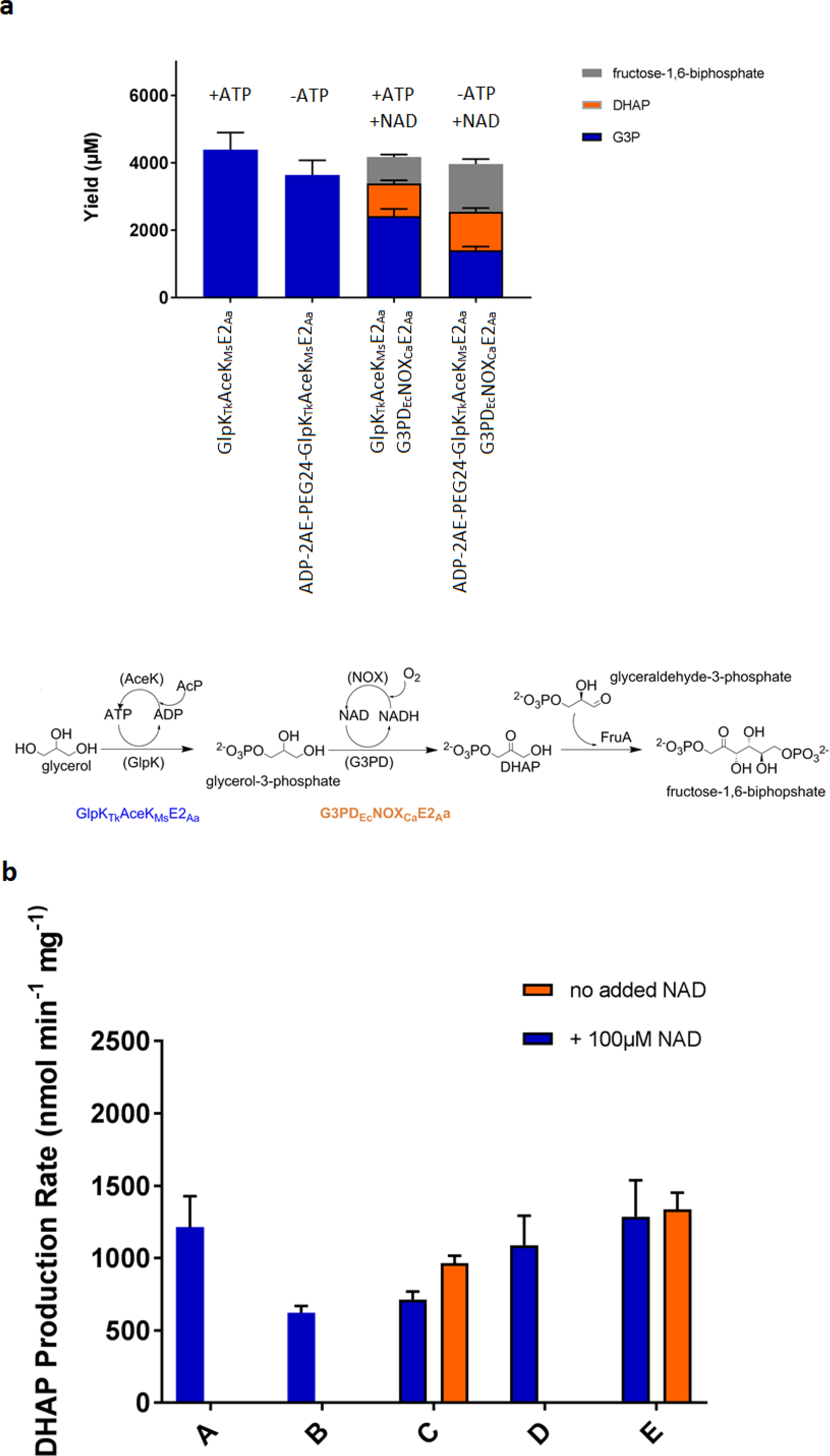
Functional tethering of modified cofactors MAL-PEG_24_-2*AE*-ADP (a) and MAL-PEG_24_-2*AE*-NAD^+^ (b) to nanomachine fusion proteins. **a,** Glycerolkinase activity with and without the addition of 100 μM ATP catalyzed by either 2*AE*-ADP-PEG_24_-MAL-GlpK_Tk_-AceK_Ms_-E2_Aa_ or by GlpK_Tk_-AceK_Ms_-E2_Aa_ was then coupled with glycerol-3-phosphate dehydrogenase activity (G3PD_Ec_-NOX_Ca_-E2_Aa_) and aldolase activity (FruA_Sc_ with glyceraldehyde-3-phosphate as added donor substrate) to demonstrate the production of fructose-1,6-biphosphate from 10 mM glycerol using 2*AE*-ADP-PEG_24_-MAL-GlpK_Tk_-AceK_Ms_-E2_Aa._ A scheme of the three step reaction involved is illustrated beneath the graph. **b,** Glycerol-3-phosphate dehydrogenase activity (DHAP production rate) using 10 mM glycerol-3-phosphate with and without the addition of 100 μM NAD^+^ catalyzed by 2*AE*-NAD^+^-PEG_24_-MAL-G3PD_Ec_-NOX_Ca_-E2_Aa_ created under different reducing and tethering conditions: untethered control G3PD_Ec_-NOX_Ca_-E2_Aa_ (A), 1 mM TCEP and 1 equivalent MAL-PEG_24_-2*AE*-NAD^+^ (B), 0.1 mM TCEP and 1 equivalent MAL-PEG_24_-2*AE*-NAD^+^ (C), no TCEP and 1 equivalent MAL-PEG_24_-2*AE*-NAD^+^ (D), 1 mM TCEP and 5 equivalent MAL-PEG_24_-2*AE*-NAD^+^ (E). All reactions were conducted for 30 minutes at 37 °C, and products analysed by LCMS as described in analytical methods.

#### Accurate mass determination of MAL-PEG_24_-2AE-NAD^+^ tethered to G3PD_Ec_-NOX_Ca_-Est2_Aa_ by LC-MS proteomics

The accurate mass of G3PD_Ec_-NAD_teth_-NOX_Ca_-Est2_Aa_ conjugates was determined by denaturing liquid chromatography–mass spectrometry (LC-MS). Protein samples were spiked with formic acid (FA) to a final concentration of 0.1% (v/v) and separated by reverse-phased liquid chromatography on an UltiMate 3000 RSLC nano system (ThermoFisher Scientific) fitted with a 50 × 4.6 mm, 5 μM particle size, 300 Å pore size PLRP-S column (Agilent). Proteins were eluted at a flow of 250 μLmin^−1^ by applying a linear 30 min gradient from 0 to 80% solvent B (mobile phase A: 0.1% (v/v) formic acid; mobile phase B: 90% (v/v) acetonitrile/0.1% (v/v) formic acid) using an Apollo II electron spray ion source coupled to a microTOF-QII mass spectrometer (Bruker). The instrument was calibrated in positive ion mode using ESI-L Low Concentration Tuning Mix (Agilent) and LC-MS raw data were processed and deconvoluted using the MaxEnt algorithm as part of Bruker Compass DataAnalysis version 4.3.

#### Sample preparation and peptide sequencing by nanoUPLC-MSMS

G3PD_Ec_-NAD_teth_-NOX_Ca_-Est2_Aa_ protein bands were manually excised from Coomassie-stained SDS-PAGE gels and subjected to manual in-gel reduction, alkylation and tryptic digestion. All gel samples were reduced with 10 mM DTT (Sigma) for 30 min, alkylated for 30 min with 50 mM iodoacetamide (Sigma) and digested with 375 ng trypsin gold (Promega) for 16 h at 37 °C. Peptides then were separated using an UltiMate 3000 RSLC nano system (ThermoFisher Scientific), utilizing a 60 min gradient on an Acclaim Pepmap 100 column (50 cm × 75 µm id with 3 µm particles). High-resolution MS/MS data was obtained on an Orbitrap Fusion Lumos Mass Spectrometer operated in data-dependent mode, automatically switching between the acquisitions of a single Orbitrap MS scan (resolution, 120,000) every 3 s and the top-20 multiply charged precursors selected for EThcD fragmentation with a resolution of 30,000 for Orbitrap MS-MS scans.

#### Mass spectra database searching

Orbitrap MS/MS data was searched against a focused decoy database containing G3PD_Ec_-NOX_Ca_-Est2_Aa_ and common contaminant protein sequences using the Byonic search engine (Protein Metrics) with tolerance of 5 ppm for precursor ions and 10 ppm for product ions. Enzyme specificity was tryptic and allowed for up to 2 missed cleavages per peptide. A Wildcard search with a range of +75 to +2200Da facilitated confident peptide identification (< 1% FDR) and spectrum counting of PEGylated cysteine residues. Variable modifications were set for NH_2_-terminal acetylation or protein N-termini, oxidation of methionine or tryptophan, and carbamidomethyl modification of cysteine.

All of the cysteine containing peptides from the tryptic digest were able to be observed by mass spectrometry in the unconjugated sample, but the conjugated sample was missing the peptide corresponding to the linker cysteine, and instead peptides corresponding to the conjugated peptide were observed (Supplementary Figure 4).

**Supplementary Figure 4:**
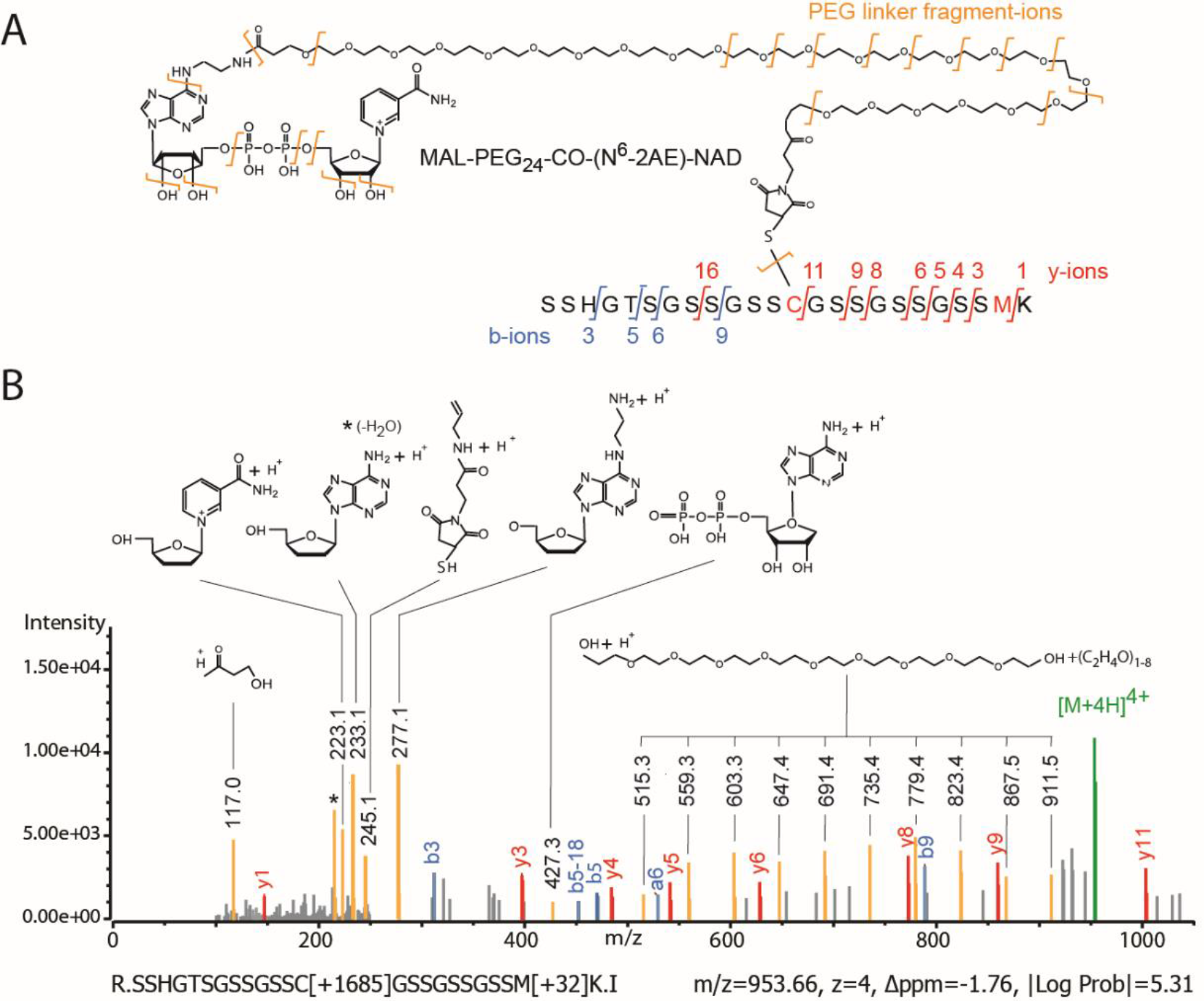
Confirmation of functional Mal-PEG_24_-2*AE*-NAD^+^ linkage to G3PD_Ec_-NOX_Ca_-Est2_Aa_ residue Cys369. **a,** Cartoon of the chemical structure of MAL-PEG_24_-2*AE*-NAD^+^ conjugated to G3PD_Ec_-NOX_Ca_-Est2_Aa_ residue Cys369 of tryptic peptide SSHGTSGSSGSSCGSSGSSGSSMK highlighting different MSMS fragment ions (peptide b-ions, blue; peptide y-ions, MAL-PEG_24_-2AE-NAD^+^ CID ions, gold). **b,** Annotated LC-MSMS evidence spectrum for a high-scoring R.SSHGTSGSSGSSC[+1685]GSSGSSGSSM[+32]K.I peptide highlighting peptide b- and y-ions (blue, red) as well as the observed masses and chemical structures of matching MAL-PEG_24_-2*AE*-NAD^+^ fragments. The observed +1685Da mass modification and fragmentation pattern is consistent with G3PD_Ec_-NOX_Ca_-Est2_Aa_ Cys396 being tethered to a functional MAL-PEG_24_-2*AE*-NAD^+^ linker.

### Nanomachine immobilization onto agarose beads and nanomachine conjugation to modified cofactors

#### Synthesis of thiohexyltrifluoroketone (hTFK)

1,6-Hexanedithiol (601 mg, 0.612 mL, 4 mmol) was added to a stirred mixture of NaHCO_3_ (4 mmol) in anhydrous dichloromethane (8 mL) under a nitrogen atmosphere. Bromotrifluoroacetone (0.415 mL, 0.4 mmol) was then added dropwise. Reaction was monitored by TLC. The reaction mixture was stirred under N_2_ for 5 days at room temperature and then poured into 50 mL water. After extraction with ether (3 × 30 mL), the organic solvent was dried (MgSO_4_) and the solvent removed under reduced pressure. Thiohexyltrifluoroketone was characterised by LC-MS; aHPLC (20 – 80% gradient MeCN into H_2_O, 0.1% TFA) gave a single peak at 6.6 min (λ = 214 nm), > 90% purity, ESI (negative scan mode) found 259.16 amu; calculated MW = 260.33. hTFK could be used directly in loading the DVS-modified beads.

#### Synthesis of hTFK-vinylsulfone activated beads

4% crosslinked agarose (Sepharose CL-4B, GE Healthcare) was functionalized by treatment with divinylsulfone (DVS) to yield the vinylsulfone decorated agarose as an aqueous slurry with vinyl sulfones at approximately 1 mmol mL^−1^ of slurry. To 500 g of damp drained Sepharose CL-4B was added 500 mL of 0.5 M Na_2_CO_3_ (pH 12) and 5000 µL divinylsulfone. The resulting suspension was stirred gently for 70 min at room temperature before being washed extensively with water. The solution was stored as a 1:1 *w*/*v* slurry in 50% ethanol/water. This activated DVS-agarose was then reacted with thiohexyltrifluoroketone (hTFK) at approximately 5 molar percent ratio for between 6 h and overnight before capping all remaining vinyl sulfone functionalities with 2-mercaptoethanol. Saturated NaHCO_3_ solution (20 mL), and thiohexyl trifluoroketone (1.2 mL ethanol solution containing 26 mg of compound, 0.1 mmol, 5% loading) were added to the divinyl sulfone resin (200 mL, 16-20 mmol, 50% slurry in 1:1 ethanol/water) and stirred at room temperature for 2 h; excessive reactive sites were blocked by the addition of beta-mercaptoethanol (2.8 mL, 40 mmol) and washed with 50% ethanol/water until no smell was evident. The hTFK-loaded agarose gel obtained was filtered, washed and stored as a 1:1 slurry in 50% ethanol/water for further use. The synthesis scheme is summarized below (Supplementary Scheme 2).

**Supplementary Scheme 2.**
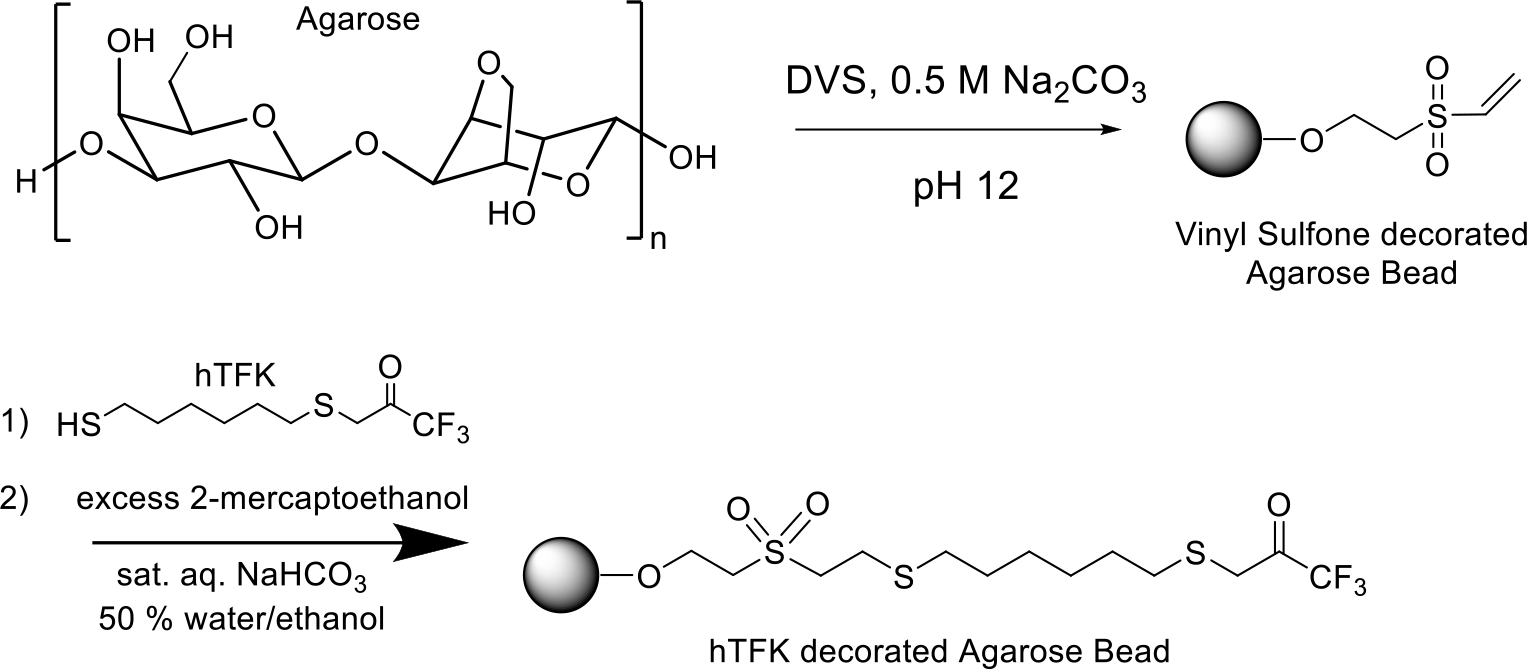
Scheme for the preparation of hTFK decorated agarose beads via vinylsulfone activation.

#### Immobilization of GlpK_Tk_-AceK_Ms_-Est2_Aa_ to Sepharose-DVS-hTFK

A lysate from 8 g GlpK_TK_-AceK_Ms_-Est2_Aa_ cells prepared as described previously (200 mL) was added to 25 g Sepharose-DVS-hTFK, and the slurry mixed at 4 °C. The loss of esterase activity in the supernatant was monitored and after 2.5 h there was no further loss of esterase activity, and the adsorbent containing 5.2 U esterase per g beads, corresponding to an estimated 0.8 mg GlpK_Tk_-AceK_Ms_-Est2_Aa_ per g, or 6 nmol GlpK_Tk_-AceK_Ms_-Est2_Aa_ per gram, was filtered and washed.

#### Tethering of MAL-PEG_24_-2AE-ADP to immobilized Sepharose-DVS-TFK-GlpK_Tk_-AceK_Ms_-Est2_Aa_

The Sepharose-DVS-GlpK_Tk_-AceK_Ms_-Est2_Aa_ (25 g) was incubated in Tris-buffered saline pH 7.0 containing 1 mM TCEP for 1.5 h at 4 °C before being washed with extensively degassed PBS containing 0.5 mM EDTA. An equal volume of this buffer was added to the slurry together with 0.8 μmol MAL-PEG_24_-2*AE*-ADP and the mixture allowed to react for 6 h at 4 °C with mixing. The slurry was then filtered and washed with TBS.

#### Immobilization of G3PD_Ec_-NOX_Ca_-Est2_Aa_ to Sepharose-DVS-hTFK

To lysate from 10.6 g G3PDEc-NOX_Ca_-Est2_Aa_ cell paste prepared as described for the purification of G3PD_Ec_-NOX_Ca_-Est2_Aa_ above was added 80 g Sepharose-DVS-hTFK and the mixture stirred gently for 100 min at 4 °C. The slurry was filtered and washed with extensively degassed PBS containing 0.5 mM EDTA and 10 μM TCEP.

#### Tethering of MAL-PEG_24_-2*AE*-NAD^+^ to immobilized Sepharose-DVS-TFK-G3PD_Ec_-NOX_Ca_-Est2_Aa_

To 35 mL of the slurry was added an equal volume of this buffer together with 580 nmol MAL-PEG_24_-2*AE*-NAD^+^ and the mixture allowed to react at 4 °C with mixing for 30 min before being filtered and washed with PBS containing 1 mM TCEP.

### Reactor Assembly

In line with the intended modular, hierarchal organization of our nanomachine technology, we optimized each nanomachine reactor individually and then combined them into a serial multi-enzyme D-fagomine nanofactory as shown in Figure 2.

#### The Phosphotransfer Reactor

For the preparation of the phosphotransfer reactor (Figure 2), 40 mg of GlpK_*Tk*_-AceK_*M*_s-Est2_*Aa*_ protein (296 nmol) was immobilized onto 25 g of Sepharose–hexyl-DVS-TFK beads. The immobilized GlpK_*Tk*_-AceK_*M*_s-Est2_*Aa*_ was treated with 0.1 mM TCEP, washed with PBS containing 0.5 mM EDTA then reacted with six equivalents MAL-PEG_24_-2*AE*-ADP for 6 h at 4 °C before being washed with reaction buffer (0.2 M sodium citrate buffer pH 7.9). The resultant immobilized cofactor-tethered nanomachine beads were analyzed for glycerol kinase activity in the presence and absence of ATP in batch reactions, and demonstrated to have ∼30% tethering efficiency (activity without added ADP calculated as the percentage of activity with added ADP). The resultant immobilized nanomachine beads were then packed into a 25 mm*15 mm Benchmark column (Kinesis, Australia) to a packed bed volume of 21.2 mL and performance assessed in a flow reactor system.

A bioreactor packed with the immobilized GlpK_*Tk*_-ATP_teth_-AceK_*M*_s-Est2_*Aa*_ nanomachine beads was found to convert 10 mM glycerol and 10 mM acetyl phosphate to G3P and acetate with approximately 60% efficiency at the optimal flow rate of 0.25 mL min^−1^ (Supplementary Figure 5a). This resulted in a space time yield of 70 mg G3P L^−1^ hr^−1^ mg^−1^ protein. The bioreactor stability was further assessed by continuing to run the phosphotransfer reactor for a total time of 870 minutes resulting in a total 14222 turnovers of the tethered cofactor (Figure 3).

#### The Oxidation Reactor

For the preparation of the G3PD_Ec_-NOX_Ca_-Est2_Aa_ oxidation reactor (step 2 in Figure 2), 80 mg of G3PDEc-NOXCa-Est2Aa protein (647 nmol; 1260 esterase U) was immobilized onto 80 g of Sepharose–hexyl-DVS-TFK. The immobilized G3PDEc-NOXCa-Est2Aa was treated with TCEP, washed with degassed, sparged PBS containing 0.5 mM EDTA then reacted with six equivalents MAL-PEG_24_-2*AE*-NAD^+^ for 6 h at 4 °C before being washed with PBS. The resultant immobilized cofactor-tethered nanomachine beads were analyzed for glycerol-3-phosphate dehydrogenase activity in the presence and absence of NAD^+^ in batch reactions, and demonstrated to have ∼ 80% tethering efficiency (activity without added NAD^+^ calculated as the percentage of activity with added NAD^+^). The resultant immobilized nanomachine beads were then packed into a 250 mm × 15 mm Benchmark column (Kinesis, Australia) to a packed bed volume of 28.3 mL and assessed in a flow reactor system.

A column packed with the adsorbent was found to convert 10 mM G3P to DHAP with about 40 – 50% efficiency at a flow rate of 0.25 mL min^−1^ (Supplementary Figure 5b). This resulted in a space time yield of 2.60 mg DHAP L^−1^ hr^−1^ mg^−1^ protein. The bioreactor stability was further assessed by continuing to run the oxidation reactor for a total time of 6000 minutes resulting in a total 1843 turnovers of the tethered cofactor (Figure 3).

#### The Aldol Addition Reactor

For the preparation of immobilized nanomachine beads for the aldol addition reactor, 20 mg of FruA_*Sc*_-Est2_*Aa*_ protein was reacted with 20 g of Sepharose–hexyl-DVS-TFK beads. The resultant immobilized aldolase nanomachine beads were then packed into a 150 mm × 15 mm Benchmark column (Kinesis, Australia) to a final depth of 10 cm (17.7 mL packed bed volume) and assessed in a flow reactor system. Optimal flow rate was assessed for the aldol reactor and found to be 0.1 mL min^−1^, with approximately 86% and 98% conversion of 5 mM *N*-Cbz-3-aminopropanal and 5 mM DHAP respectively, under these conditions (Supplementary Figure 5c).

**Supplementary Figure 5.**
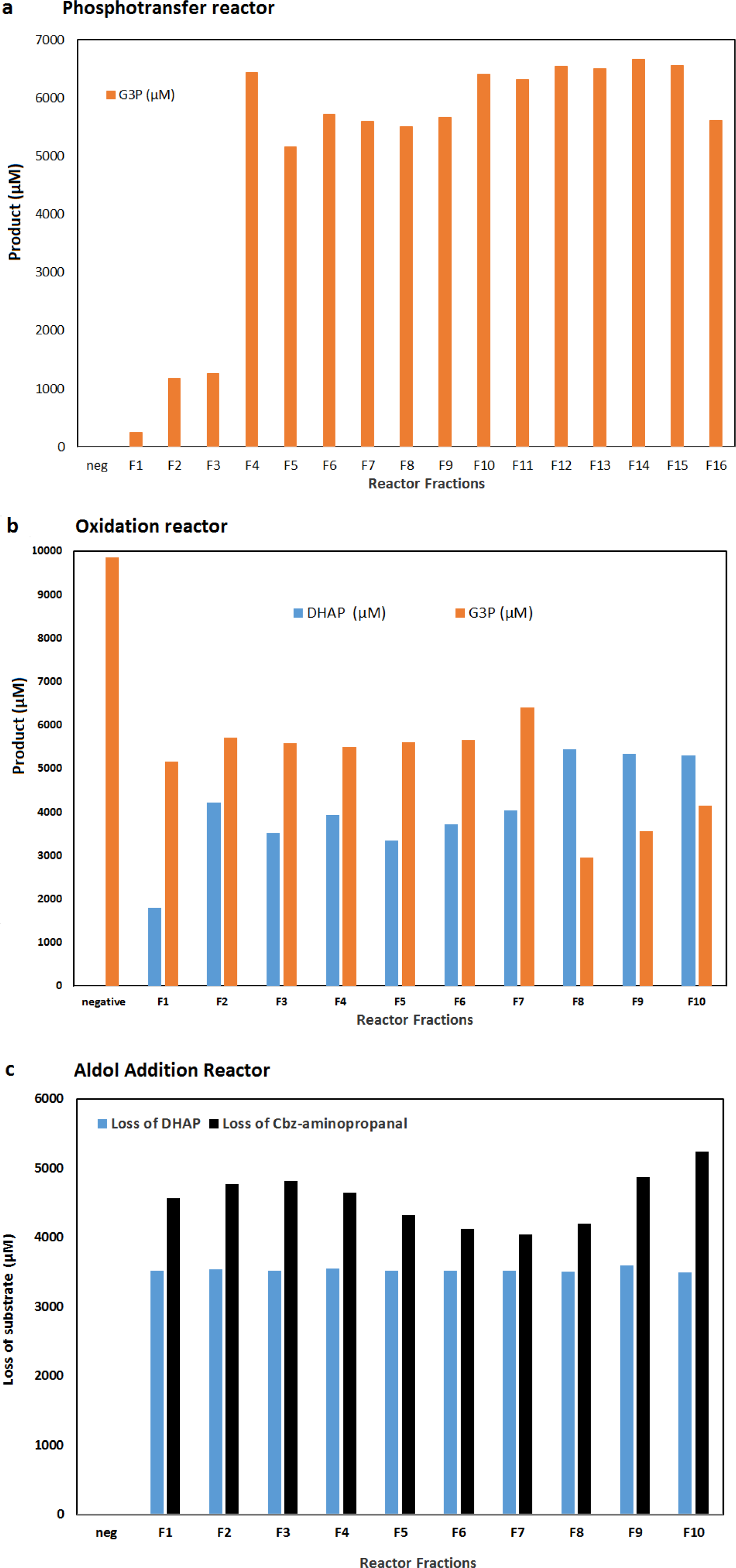
Assembly and testing of each of the nanomachine reactors used to assemble the Nanofactory. **a,** Phosphotransfer Reactor: Conversion of glycerol and acetyl phosphate (10 mM each) to G3P and acetate by immobilized GlpK_Tk_-ATP_teth_-AceK_Ms_-Est2_Aa_ in a packed bed reactor column (1.5 cm id, 12 cm) run at a flow rate of 0.25 mL min^−1^, as determined by LCMS analysis of 5 mL fractions. **b,** Oxidation reactor: conversion of G3P to DHAP in a flow reactor. The immobilized G3PD_Ec_-NAD_teth_-NOX_Ca_-Est2_Aa_ nanomachine beads were used to prepare a packed bed reactor column (1.5 cm id × 16.5 cm). 10 mM G3P pH 8 was passed through the column at a flow rate of 0.25 mL min^−1^ and the amount of G3P remaining and DHAP produced determined by LCMS for 5 mL fractions F1 to F10. **c,** Aldol addition reactor with Fru_Sc_-E2_Aa_: conversion of Cbz-aldehyde and DHAP into *N-*Cbz-3*S*,4*R*-ADHOP in a flow reactor. The immobilized Fru_Sc_-E2_Aa_ nanomachine beads prepared in the presence of 10 μM TCEP were used to prepare a packed bed reactor column (1.5 cm id × 16.5 cm). 5 mM *N*-Cbz-3-aminopropanal and DHAP in 50 mM citrate buffer pH 7 was passed through the column at a flow rate of 0.1 mL min^−1^ and the amount of DHAP and *N*-Cbz-3-aminopropanal remaining quantified by LCMS for 5 mL fractions F1 to F10.

### Enzyme activity assays (In batch and in flow)

#### Glycerol kinase activity

Glycerol kinase assays were performed at room temperature in 1 mL volume with direct detection of ADP and ATP by HPLC analysis of reaction supernatant. A typical reaction contained 1 mM glycerol, 10 mM MgCl_2,_ 50 mM NaHCO_3_ buffer pH 9.0, 1 mM ATP with approximately 2 µg mL^−1^ enzyme (∼35 nM). Kinetics were determined by varying the concentrations of ATP or glycerol whilst maintaining the other in excess, and kinetic determinants calculated using Hyper™ (J.S. Easterby, Liverpool University) or GraphPad Prism (GraphPad Software Inc., USA). Substrate and cofactor concentrations ranged from 0.1 to 10 × *K*_M_.

#### Acetate kinase activity

Acetate kinase assays were conducted in the same manner as the glycerol kinase assays described above, replacing ATP with ADP, and glycerol with acetyl phosphate or phosphoenol pyruvate. Kinetics were determined by varying the concentrations of ADP or acetyl phosphate or phosphoenol pyruvate whilst maintaining the other components in excess, and kinetic determinants calculated using Hyper (J.S. Easterby, Liverpool University). Substrate and cofactor concentrations ranged from 0.1 to 10 × *K*_M_.

#### Glycerol-3-phosphate dehydrogenase activity

Glycerol-3-phosphate (G3P) dehydrogenase activity was determined from the oxidation of glycerol-3-phosphate to DHAP in 50 mM sodium phosphate pH 9.0 for individual enzyme assays, or at pH 8.0 for combined multienzyme reactions, with the reaction progress followed by monitoring the production of NADH spectroscopically at 340 nM (ε_340 nm_ 6.22 mM^−1^ cm^−1^), or by direction detection of both G3P (substrate) and DHAP (product) using LCMS (see Analytical methods and Supplementary Figure 6), with one unit of glycerol-3-phosphate activity defined as the amount required to oxidize 1 μmol G3P in one minute at ambient temperature. Kinetics were determined by varying the concentration of G3P and NAD^+^ from 0.1 to 10 × *K*_M_ and kinetic determinants were calculated using Hyper™ (J.S. Easterby, Liverpool University) or GraphPad Prism (GraphPad Software Inc., USA).

#### NADH oxidase activity (untethered)

NADH oxidase activity was determined from the oxidation of 0.1 mM NADH in 50 mM sodium phosphate pH 7 containing 1 mg mL^−1^ BSA, with the loss of NADH monitored spectroscopically at 340 nM (ε_340 nm_ 6.22 mM^−1^ cm^−1^), with one unit of NADH oxidase activity defined as the amount required to oxidize 1 μmol NADH in one minute at ambient temperature. Kinetics were determined by varying the concentration of NADH from 0.1 to 10 × *K*_M_ and kinetic determinants were calculated using Hyper™ (J.S. Easterby, Liverpool University) or GraphPad Prism (GraphPad Software Inc., USA).

#### Esterase activity

Esterase activity was determined from the hydrolysis of *p*-nitrophenyl acetate (Sigma) in 50 mM sodium phosphate pH 7 containing 1 mg mL^−1^ BSA and with a typical reaction containing 0.4 mM *p*-nitrophenyl acetate. The hydrolysis of *p*-nitrophenyl acetate was determined spectroscopically by the increase in absorbance at 405 nm due to production of *p*-nitrophenol (ε_405 nm_ 18 mM^−1^ cm^−1^), with one unit of esterase activity defined as the amount required to hydrolyze 1 μmol *p*-nitrophenyl acetate in one minute at ambient temperature. Kinetics were determined by varying the concentration of substrate from 0.1 to 10 × *K*_M_ and kinetic determinants were calculated using Hyper™ (J.S. Easterby, Liverpool University) or GraphPad Prism (GraphPad Software Inc., USA).

### Analytical Methods

#### HPLC separation of ATP and ADP

HPLC separation was conducted using an Agilent Eclipse XDB column (50 mm × 4.6 mm) with isocratic elution using 75% solvent A and 25% solvent B. Solvent A: 20 mM tetrabutylammonium phosphate (TBAP) in 10 mM ammonium phosphate buffer pH 4.0; solvent B: acetonitrile. Flow rate 1 mL per minute, detection at 240 nm using diode array detector (Agilent Technologies, USA). Peaks eluted at the following retention times: ADP 1.2 min, ATP 1.8 min.

#### LCMS analysis of glycerol-3-phosphate (G3P), dihydroxyacetone phosphate (DHAP) and aldol products

G3P and DHAP were separated using a modification of the method described in Prieto-Blanc *et al*., 2010 ^32^. Chromatographic conditions were SIELC ObeliscN column (100 mm × 2.1 mm) with isocratic elution using 20% mobile phase A, 80% mobile phase B for 5 minutes. Mobile phase A: 25 mM ammonium formate pH 4.0; mobile phase B: acetonitrile. Mass spectrophotometric detection was conducted using API-ES negative mode with an Agilent 6120 Quadropole LCMS. Compounds were qualitatively detected by ion scanning in both positive and negative mode using know standards, and then quantified based on selected ion monitoring (SIM) monitoring of relevant ions. Glycerol-3-phosphate was quantified by selected ion monitoring of ion [M]^−^ *m/z* = 171.06, DHAP quantified by selected ion monitoring of ion [M]^−^ *m/z* = 169.04, after establishing suitable selected ions using positive and negative scanning of standards. Quantitation was based on comparison to standard calibration curves produced in the same manner. *N*-Cbz-3-aminopropanal and *N*-Cbz-3*S*,4*R-*ADHOP were detected and quantified using absorbance A_214nm_ and selected ion monitoring of ion [M-H]^−^ *m/z* = 376.09 (*N*-Cbz-3*S*,4*R-*ADHOP) by comparison with calibration curves made using a synthesised standard (Supplementary Figure 6). The *N*-Cbz-3*S*,4*R-*ADHOP standard was synthesised from DHAP (Sigma-Aldrich) and *N*-Cbz-3-aminopropanal (Sigma-Aldrich) using enzymatic aldol addition with purified FruA_Sc_ and the resultant *N*-Cbz-3*S*,4*R-*ADHOP was purified essentially as described by Castillo and colleagues ^17^. ^1^H-NMR spectroscopy confirmed the purity of the standard (Supplementary Figure 6c) and this standard was then used to identify and quantitate the *N*-Cbz-3*S*,4*R-*ADHOP produced from the nanofactory by selected ion monitoring.

**Supplementary Figure 6.**
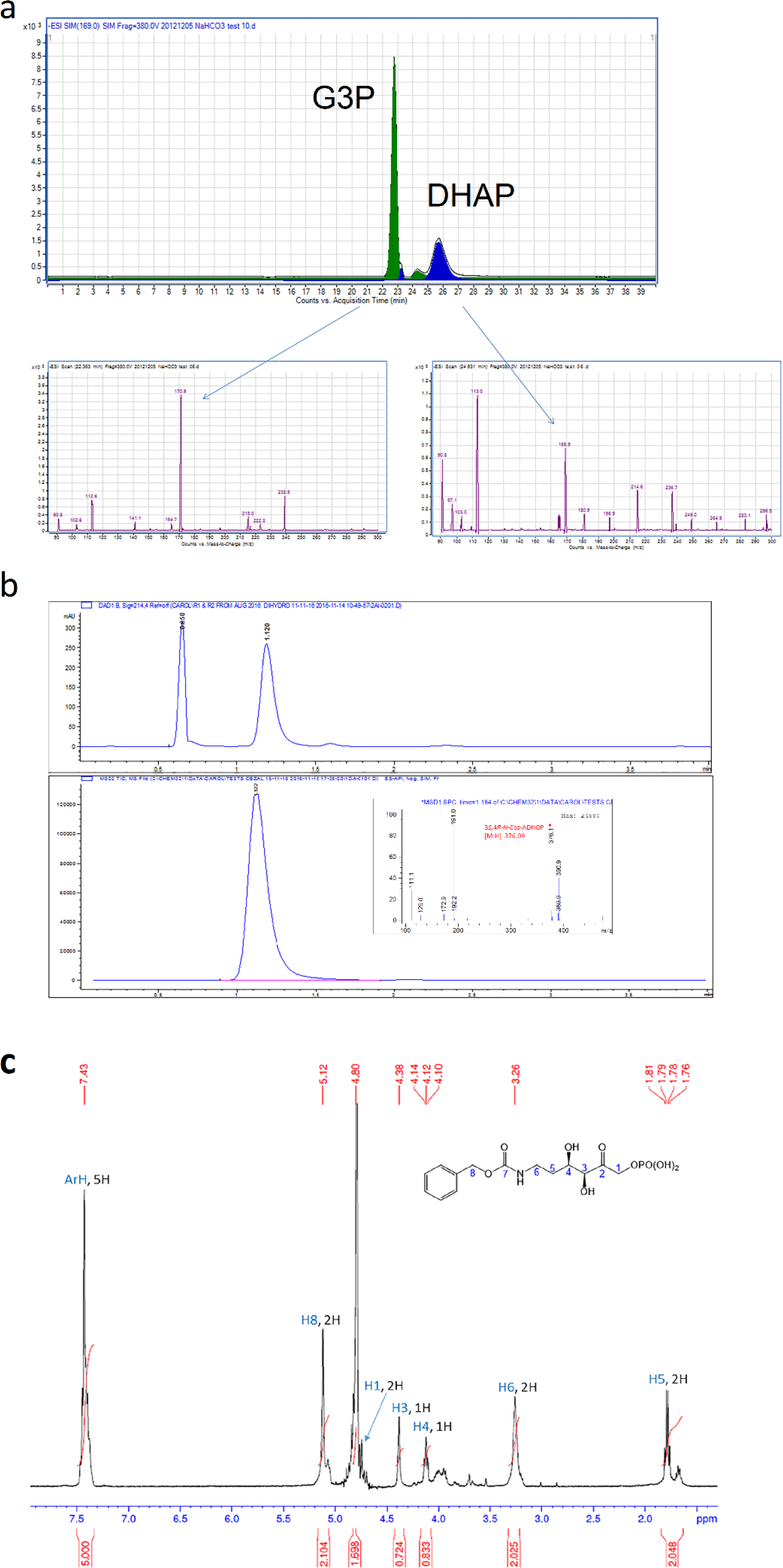
HPLC and HPLC-MS traces and spectra for all components quantified to measure the conversion of glycerol and aldehydes into chiral aldol products. **a,** HPLC separation and mass spectrometry identification of glycerol-3-phosphate (G3P) and dihydroxy acetone phosphate (DHAP). The dominant selected ions identified here (m/z 171^−^ for G3P and m/z 169^−^ for DHAP) were then used for SIM analyzes of subsequent reactions. **b,** HPLC separation of *N*-Cbz-3-aminopropanal and *N-*Cbz-3*S*,4*R*-ADHOP showing absorbance A_214nm_ (upper panel) and mass spectrometry identification of *N-*Cbz-3*S*,4*R*-ADHOP (inset). **c**, the authenticity of *N-*Cbz-3*S*,4*R*-amino-3,4-dihydroxy-2-oxyhexyl phosphate (*N*-Cbz-3*S*,4*R*-ADHOP) produced by the nanofactory was confirmed by ^1^H-NMR analysis of *N*-Cbz-3*S*,4*R*-ADHOP prepared by reaction of commercially available *N*-Cbz-3-aminopropanal and DHAP catalysed by FruA, and isolated by HPLC

